# Knowledge Inclusive Machine Learning for Disease Gene Prioritisation

**DOI:** 10.64898/2026.04.29.721522

**Authors:** Chathura J. Gamage, Yu Xia, Ravisha Rupasinghe, Sachith Seneviratne, Damith Senanayake, Tamasha Malepathirana, Asela Hevapathige, Mark Corbett, Terence J. O’Brien, Steven Petrou, Samuel F. Berkovic, Ingrid E. Scheffer, Jozef Gecz, Melanie Bahlo, Mark F. Bennett, Saman Halgamuge

## Abstract

The predictive performance of machine learning models depends on the context available to them. In disease gene prioritisation, this context comprises two forms: specific context from sample-level experimental data, such as gene expression and protein–protein interaction networks, and general context from accumulated and curated biological knowledge capturing established relationships among genes, diseases, and pathways. Neither is sufficient alone: experimental data are sensitive to dataset-specific noise and lack broader biological grounding, while curated knowledge lacks the resolution required for gene-level discrimination. Consequently, most machine learning approaches relying solely on experimental data risk learning spurious correlations rather than underlying biology. Here we introduce Knowledge Inclusive Machine Learning (KIML), a paradigm that integrates both context types within a unified analytical pipeline. KIML combines experimental data with two types of general context: literature-derived representations from PubMed and structured biomedical knowledge graphs. We evaluate the approach on Developmental and Epileptic Encephalopathy and benchmark it against recent methods using publicly available datasets. Performance is assessed using temporal-split evaluation and biological evaluations, including ontology enrichment analysis. KIML consistently outperforms existing approaches, providing improved predictive accuracy and biologically meaningful insights. Furthermore, the framework generates interpretable explanations of gene prioritisation and demonstrates strong generalisability across six additional diseases.

## 1 Introduction

Up to 80% of individuals with rare diseases remain undiagnosed following genomic sequencing, often because the underlying pathogenic variants occur in genes not yet associated with disease [1]. Computational identification of such genes, commonly referred to as disease gene prioritisation, is therefore critical for improving diagnostic rate and enabling the development of targeted therapies [2]. Laboratory-based experiments to determine disease causality are time-consuming and costly, and are therefore only feasible for a small number of candidate genes. Computational approaches can substantially narrow this search space, guiding experimental efforts towards the most promising genes.

Machine Learning (ML) is becoming an increasingly important approach in disease gene prioritisation [3, 4], as it can identify complex non-linear patterns that are difficult to detect through traditional approaches. This has shown the potential to narrow down candidate gene lists, reduce the search space for laboratory-based experiments, and accelerate the pace of gene discovery. However, the predictive power of any ML model is fundamentally bounded by the context it has access to during training. In disease gene prioritisation, we categorise the available context into two forms. The first is specific context: it refers to the experimental data, such as gene expression profiles and protein-protein interaction (PPI) measurements [5, 6]. Such experimental data are used in their original form or with only minimal preprocessing. This type of context typically operates at the sample-level, capturing condition-specific signals within a given cohort or tissue, and providing sufficient resolution to distinguish between individual genes. The second is general context that comprises accumulated biological knowledge over many decades, including curated gene–disease associations, ontologies, scientific models of the disease, clinical insights, and structured representations such as knowledge graphs derived from the literature [7]. This type of context encodes what the scientific community already understands about a gene, a disease, and the biological processes that connect them. These two forms of context are not interchangeable, and neither is sufficient on its own. Specific context used in isolation gives a model high resolution but no prior. Without the constraints imposed by existing knowledge about the disease, ML models are prone to learning spurious correlations arising from noise and the incomplete coverage of any single experimental dataset. General context used in isolation provides those constraints, but lacks the signal at the sample level that is needed to discriminate between candidate genes in a given study. Combining the two allows the general context to act as a scientific prior, while the specific context contributes the resolution needed to refine what is currently documented.

Existing ML methods for disease gene prioritisation rely almost exclusively on experimental data (i.e., specific context) [3, 4]. To address this limitation, we propose a knowledge-inclusive Machine Learning paradigm (KIML) that expands the inputs of current ML to include any available context (both specific and general information) (Fig. 1). In this paper, our KIML pipeline integrates two forms of expert-curated knowledge, which serve as representations of general context. First, we propose to integrate PubMed literature-derived gene descriptions [8]. Advances in machine learning, particularly large language models (LLMs) [9], enable the effective transformation of unstructured biomedical text into informative embeddings that can be combined with experimental data. Therefore, we use a pretrained LLM [10, 11] to process gene descriptions from the PubMed literature and obtain text embeddings as input of expert-curated knowledge for the KIML pipeline. Second, for ML architectures including Graph Neural Networks (GNNs), which are capable of capturing relational information between genes, we utilise a Knowledge Graph [7] as input. The Knowledge Graph integrates diverse biomedical data (genes, diseases, and treatments) into a structured, interconnected format to represent human-curated gene-gene relationships. Together, these components enable KIML to incorporate both the latest scientific knowledge and established biological insights, enhancing the robustness and interpretability of gene prioritisation.

**Fig. 1:**
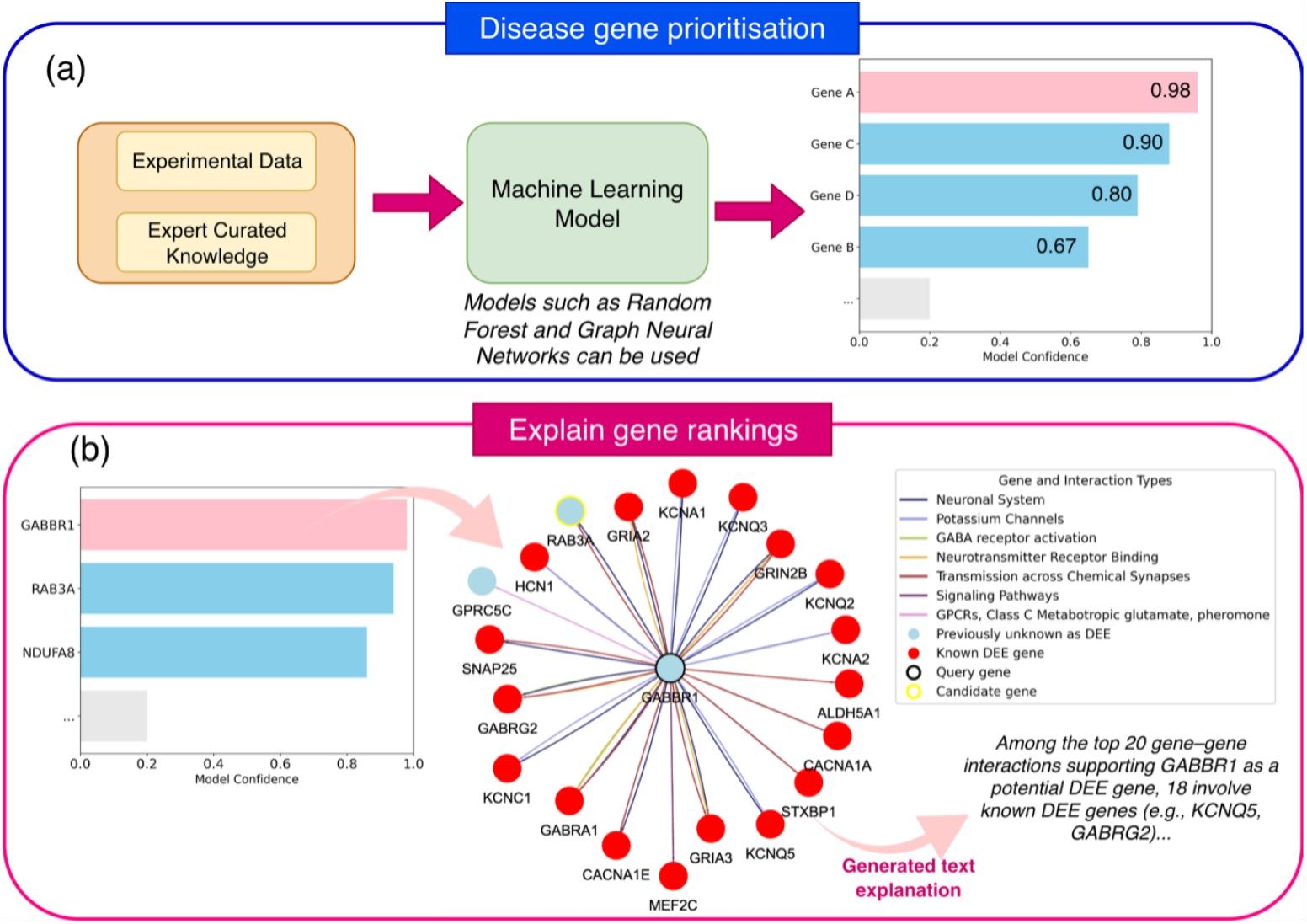
(a) Overview of a Knowledge-inclusive ML gene prioritisation pipeline. The model outputs a ranked gene list with confidence scores for disease relevance, supporting candidate gene discovery. (b) Network-based explanations link ranked candidate genes to known disease genes through biologically interpretable pathways and interaction types.

To evaluate the gene prioritisation performance of KIML, we introduce a temporal-split evaluation protocol that reflects the time-dependent nature of gene discovery. The validity of predicted disease–gene associations can only be confirmed through future discoveries, which may take years and remain incomplete. Some potential disease genes may never be experimentally investigated. Consequently, evaluation must account for the evolving state of biological knowledge. Existing methods [3, 12] typically employ random train–test splits, disregarding this temporal dependency, which can lead to unintended information overlap between training and testing data, thereby inflating performance estimates. Our protocol addresses this limitation by partitioning training and testing sets for both specific and general context using a user-defined cutoff date. Models are trained exclusively on knowledge available prior to this date and evaluated against disease genes validated subsequently, providing a more realistic and unbiased assessment. In addition, we perform biological evaluation using loss-of-function intolerance, literature-based evidence, phenotype enrichment [13], and gene ontology enrichment analyses [14].

To enable researchers to understand why certain genes are ranked highly, we employ Explainable AI (XAI) [15] to make the gene prioritisation process transparent, allowing them to validate predictions grounded on existing biological knowledge, such as known pathway connections [7]. Our explainable results are integrated into a gene prioritisation web interface, which displays an interactive network of biological relationships (e.g., pathways derived from the Knowledge Graph) responsible for the prediction of each disease gene. Such explanations can also yield insights into the biology of novel genes, as they can highlight biological mechanisms through pathway contributions and neighbouring genes. Our code and an interactive web interface for exploring results and explanations are publicly available at: https://dee-list-hz53fjmjutnybiqjb2iunv.streamlit.app/.

In summary, we propose KIML, a paradigm that systematically incorporates expert-curated knowledge into ML methods and develop a pipeline for disease gene prioritisation that leverages both curated knowledge and biological experimental data. We customise and test this pipeline for Developmental and Epileptic Encephalopathy (DEE) and further evaluate its disease-agnostic capability across six additional diseases. We introduce a temporal evaluation protocol as a best practice for fairly evaluating and comparing disease gene prioritisation methods, complemented by established biological evaluation approaches, demonstrating significant improvements over existing methods. Finally, we present a web interface that displays gene rankings and their corresponding explanations obtained through an Explainable AI method, enabling researchers to interpret our prioritisation results.

## 2 Results

In this section, we demonstrate the value of knowledge-inclusive machine learning through the results of integrating experimental data (specific context) with expert-curated knowledge (general context). The proposed KIML methods demonstrate generalised performance across multiple diseases: the model was trained for DEE using a specific input configuration, and the same configuration was applied to all other diseases. Additionally, we propose a novel protocol for evaluation, which improves the reliability of all disease gene prioritisation methods.

### 2.1 Proposed framework

In our KIML paradigm, we incorporate expert-curated knowledge through two representations: literature-derived gene descriptions and biological knowledge graphs. Specifically, gene descriptions are constructed by collecting titles and abstracts from PubMed publications that reference each gene. In this work, we use article titles and abstracts as the source text for constructing gene descriptions (while the approach is not limited to these sections, and other parts of the literature could also be used). These descriptions are then converted into numerical embeddings using pretrained language transformers or large language models (LLMs) [10, 11], which transform unstructured text into vector representations that can be processed by ML models (See Section 4 for details). For the knowledge graph component, we utilise HetioNet [7], which contains gene-gene relationships from 29 heterogeneous biological data sources, including pathway information and phenotypic associations.

The KIML paradigm is designed to be applicable across different ML architectures for disease gene prioritisation. However, the capacity to process different data types varies across model architectures. For instance, Random Forests (RF) can only process features of individual genes (gene-level information), while GNNs can also use relationships between genes (interaction-level information) [16]. To make the proposed pipeline compatible with any architecture, we further categorise data sources (expert-curated knowledge and experimental data) based on the type of information they provide (as shown in Fig. 2): (a) gene level data, capturing properties of individual genes such as gene expression data, and (b) interaction level data, capturing relationships between genes such as gene co-expression networks. Based on this categorisation, for architectures that can only use gene-level data, such as RF, we integrate only gene-level data from expert-curated knowledge and experimental data (Types A and C), resulting in the **Knowledge Inclusive Random Forest (KI-RF)**. For architectures that have the capability of utilising both gene and interaction level information, such as GNNs, we are able to incorporate all four data types (A, B, C, D), resulting in the **Knowledge Inclusive Graph Neural Network (KI-GNN)**. An overview of an example input configuration of the proposed KI-GNN is shown in Fig. 3.

**Fig. 2:**
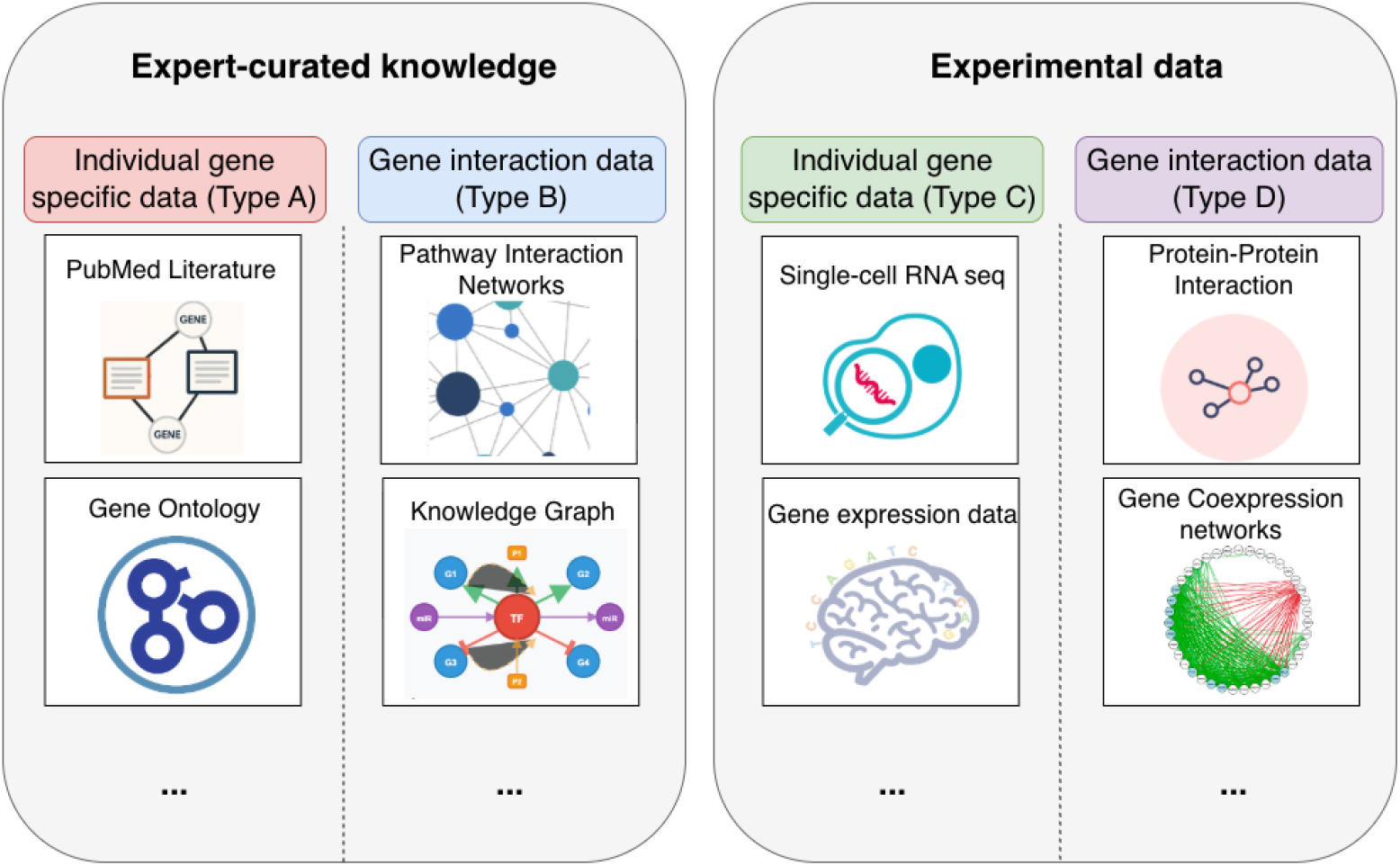
Overview of the biological data that could be used for gene prioritisation. The left panel shows expert-curated knowledge and information, and the right panel shows experimental data. Both panels have two subcategories: (1) Gene level data: individual gene-specific data (Type A and Type C) and (2) Interaction level data: the interaction data involving pairs of genes/proteins (Type B and Type D).

**Fig. 3:**
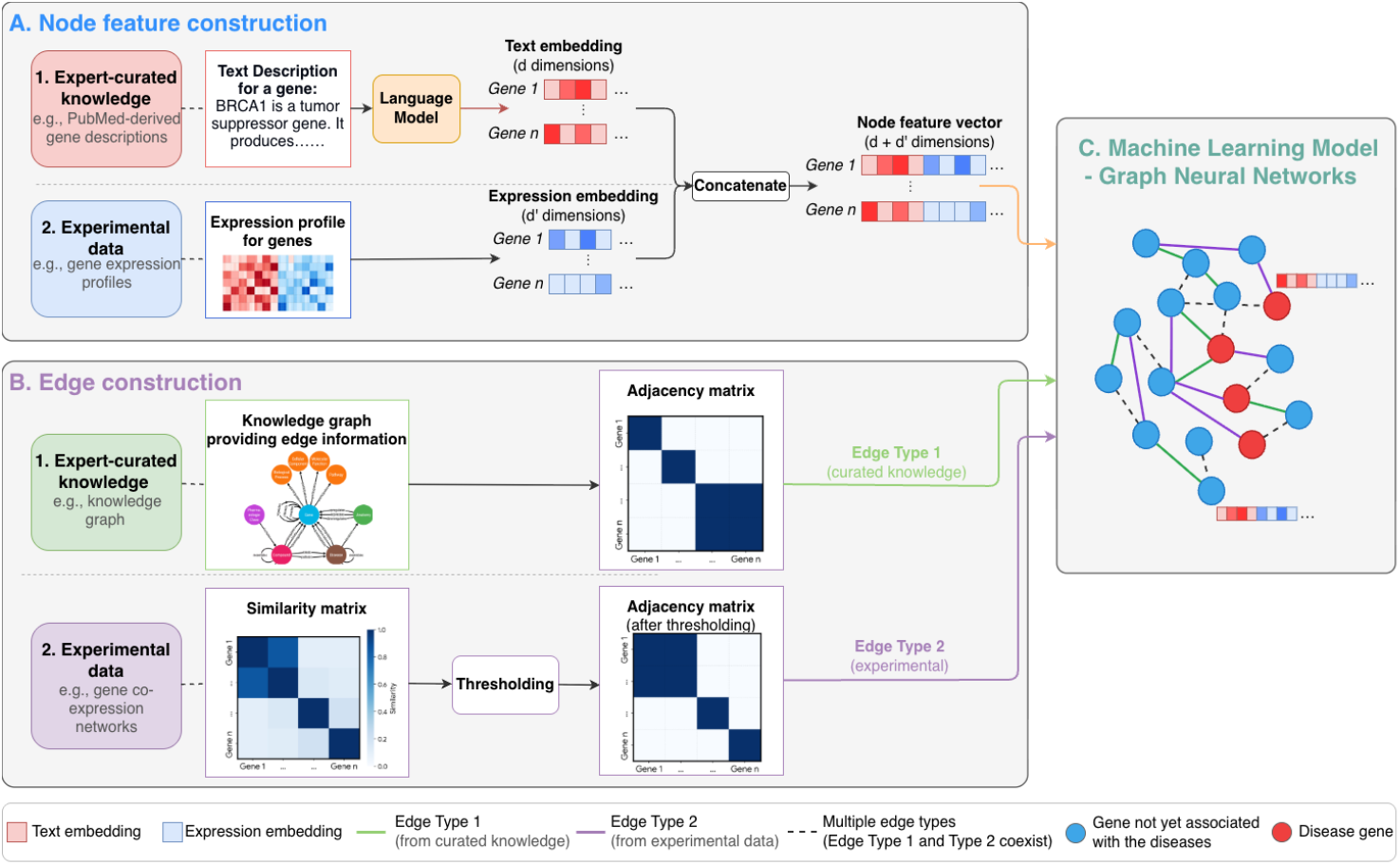
Overview of the proposed KI-GNN pipeline for disease gene prioritisation. Expert-curated knowledge and experimental data are used as input to the ML model. The illustrated pipeline serves as an example. Data sources and their combinations can be customised according to different applications.

### 2.2 Evaluation of the proposed paradigm

We evaluated gene prioritisation across three scenarios. First, we prioritised genes for Developmental and Epileptic Encephalopathy (DEE), as the primary disease of interest. Second, we focused on prioritising autosomal dominant (AD) and autosomal recessive (AR) DEE genes separately, as distinguishing inheritance-specific patterns is clinically important for understanding disease mechanisms and improving diagnosis. Third, we applied our gene prioritisation pipeline, which was optimised for DEE genes in terms of input data selection, to four additional neurological diseases and two other diseases, thereby assessing the broader applicability and generalisation capability of the methods. Although we specialise our pipeline for DEE, we empirically show that the features selected for DEE have the capability to conduct disease gene prioritisation of several other diseases. This provides confidence that the method would perform better if customised on an individual basis for those conditions, however our focus in this work remains on evaluating the feature set learned for DEE.

We compared six methods organised into three categories based on their input data sources. RF and GNN are existing methods that rely solely on experimental data, implemented using the Mantis [12] and SPEOS [3] frameworks, respectively, each with its own distinct experimental datasets. C-RF and C-GNN use the same model architectures but are trained exclusively on expert-curated knowledge, i.e., ignoring all experimental data. KI-RF and KI-GNN are our proposed Knowledge Inclusive methods that integrate both experimental data and expert-curated knowledge, extending RF and GNN through the KIML pipeline.

The input datasets used for each method are summarised in Supplementary Table 1. RF and KI-RF share the same experimental data, as do GNN and KI-GNN, providing a fair basis for evaluating the performance improvements introduced by the KIML pipeline. C-RF and C-GNN share the same expert-curated knowledge as KI-RF and KI-GNN, allowing direct assessment of the value of combining both data sources. The Mantis framework integrates experimental data from diverse public resources, including genetics databases (e.g., gnomAD, ExAC), genic intolerance scores (e.g., RVIS and MTR), functional annotations (e.g., GOA, MSigDB), gene expression and QTL datasets, GWAS resources, and disease-specific databases (e.g., CKDdb). The SPEOS framework integrates Allen Human Brain Dataset gene expression data, protein-protein interaction data from BioGRID, and single-cell RNA data. We also evaluated each method on alternative experimental datasets, with the results reported in Supplementary Table 3. Alternative experimental datasets resulted in suboptimal results compared to the original experimental datasets used in the respective methods.

#### 2.2.1 Temporal-Split Evaluation Protocol

We propose a temporal-split evaluation protocol (Fig. 4) to evaluate the gene prioritisation performances. Unlike existing gene prioritisation work, this evaluation accounts for the time-dependent nature of laboratory-based disease-gene discovery and prevents information leakage between training and testing data (as noted in the Introduction). First, a partition date is defined, and only data and expert-curated knowledge available before this date are used for training the models. In our work, this data partition date was set to 30th September 2022 (see Section 4 for the rationale and further details). After training, the models rank genes that were previously undiscovered (before the partition date), to predict which are most likely to be disease genes (blue and hatched area in Fig. 4a). Predictive performance is assessed based on how highly the models ranked the disease genes that were experimentally discovered after the partition date (blue area in Fig. 4a). The median and mean ranks of disease genes discovered after the partition date are used as an indicator of predictive performance, with lower ranks indicating better performance. We report the median rank for each method in Table 1, due to the long-tailed nature of the gene ranking distributions (Fig. 5) along with statistical significance of the observed improvements. We also evaluated the models using precision and recall at K metrics (Supplementary Table 4), which show trends consistent with the median rank results.

**Fig. 4:**
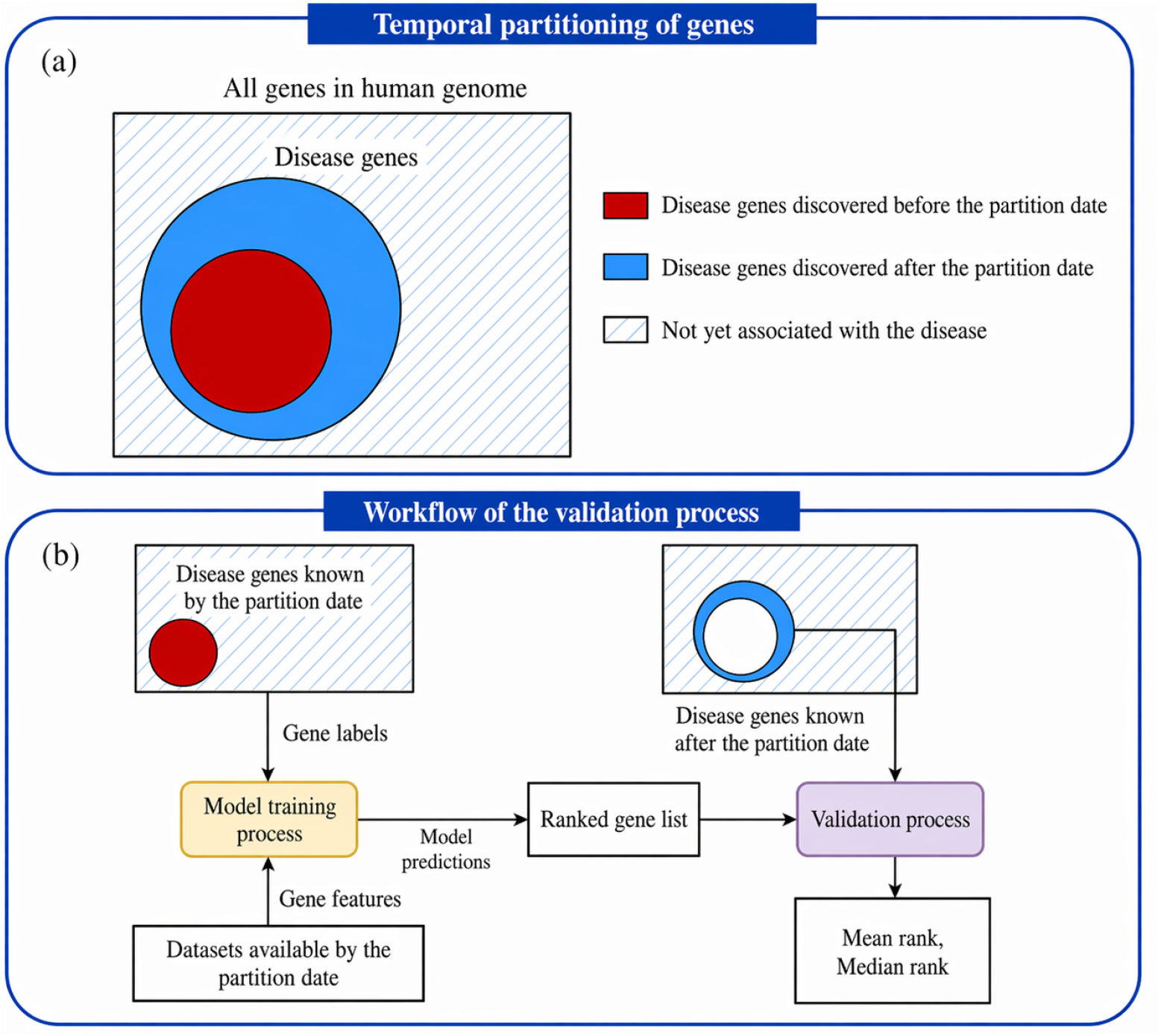
Temporal-split evaluation protocol. (a) Partition of all genes into disease genes known before the partition date (red), disease genes experimentally discovered after the partition date (blue), and those not yet associated with the disease (hatched). (b) Workflow of the evaluation process, where data prior to the partition date are used for training (red), and predictions are validated using disease genes discovered after the partition date (blue). Prediction performance is quantified using the mean and median ranks of newly discovered disease genes.

**Table 1:**
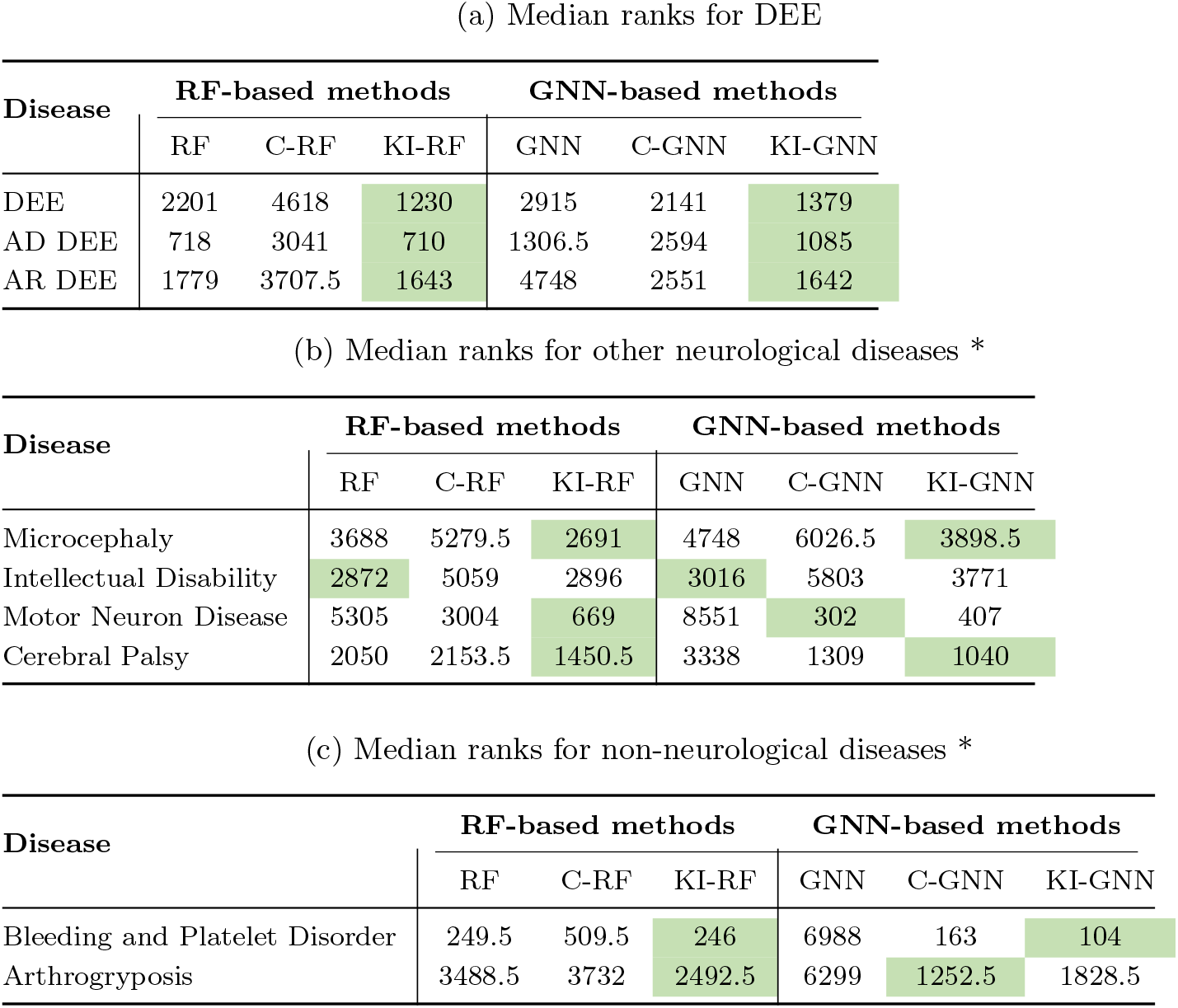
Median rank comparison across diseases. Green colour indicates the best (lowest) median rank within each method family (RF-based or GNN-based). * indicates diseases not used for model calibration.

| Disease | RF-based methods |  |  | GNN-based methods |  |  |
| --- | --- | --- | --- | --- | --- | --- |
|  | RF | C-RF | KI-RF | GNN | C-GNN | KI-GNN |
| DEE | 2201 | 4618 | 1230 | 2915 | 2141 | 1379 |
| AD DEE | 718 | 3041 | 710 | 1306.5 | 2594 | 1085 |
| AR DEE | 1779 | 3707.5 | 1643 | 4748 | 2551 | 1642 |

(b) Median ranks for other neurological diseases \*
| Disease | RF-based methods |  |  | GNN-based methods |  |  |
| --- | --- | --- | --- | --- | --- | --- |
|  | RF | C-RF | KI-RF | GNN | C-GNN | KI-GNN |
| Microcephaly | 3688 | 5279.5 | 2691 | 4748 | 6026.5 | 3898.5 |
| Intellectual Disability | 2872 | 5059 | 2896 | 3016 | 5803 | 3771 |
| Motor Neuron Disease | 5305 | 3004 | 669 | 8551 | 302 | 407 |
| Cerebral Palsy | 2050 | 2153.5 | 1450.5 | 3338 | 1309 | 1040 |

| Disease | RF-based methods |  |  | GNN-based methods |  |  |
| --- | --- | --- | --- | --- | --- | --- |
|  | RF | C-RF | KI-RF | GNN | C-GNN | KI-GNN |
| Bleeding and Platelet Disorder | 249.5 | 509.5 | 246 | 6988 | 163 | 104 |
| Arthrogryposis | 3488.5 | 3732 | 2492.5 | 6299 | 1252.5 | 1828.5 |

**Fig. 5:**
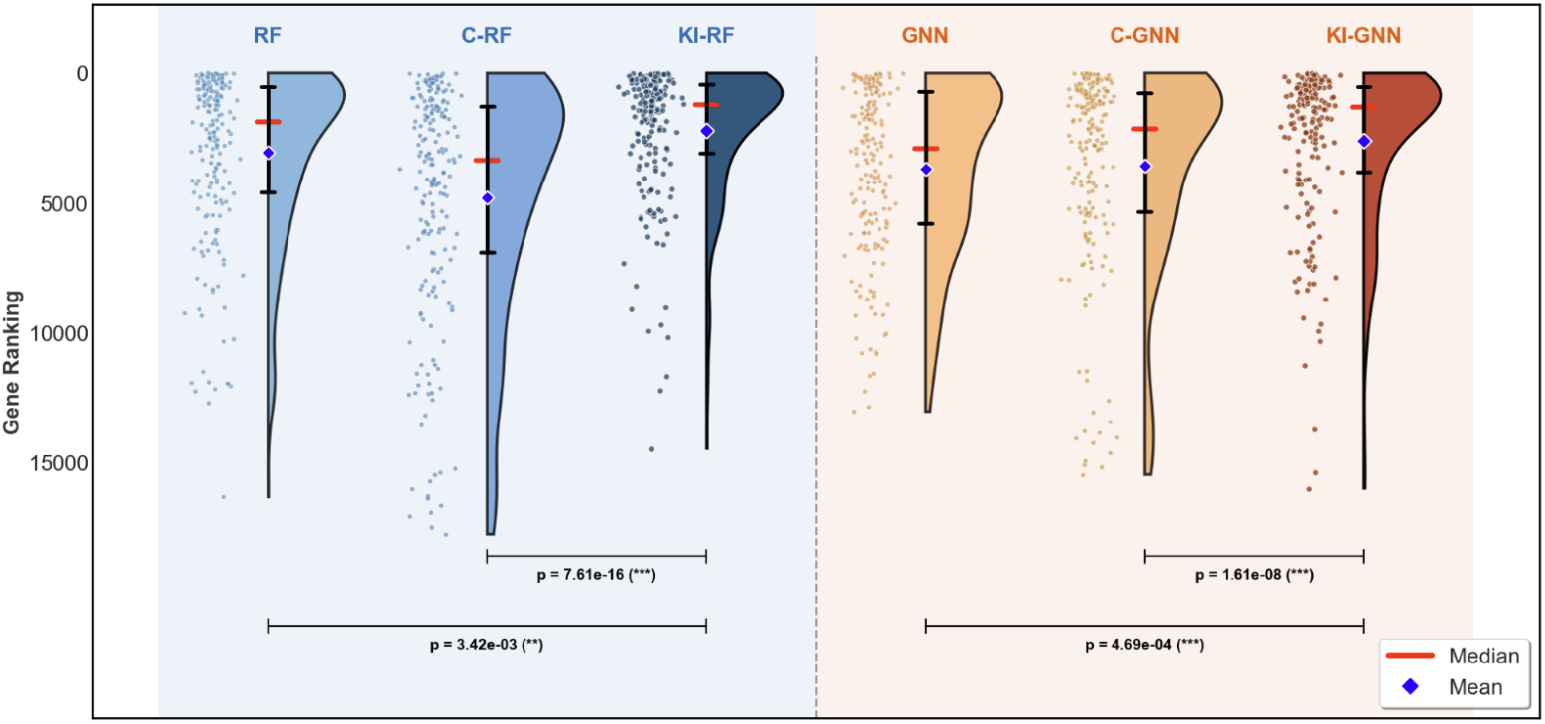
Ranking performance of each model on DEE genes discovered after the partition date. Pairwise differences were assessed using the Wilcoxon signed-rank test; significance levels are indicated below the plot (*** *p <* 0.001, ** *p <* 0.01, * *p <* 0.05, ns = not significant).

Table 1a reports median ranks for DEE, AD DEE, and AR DEE. Fig. 5 illustrates the ranking performance of each model on DEE genes discovered after the partition date (blue area in Fig. 4a). Due to the fixed partition date, the number of genes in the train-test split varies greatly between different diseases, explaining some of the variance in performance observed. The Knowledge Inclusive models consistently achieve the best median ranks across all DEE subtypes, with KI-RF improving from 2201 (RF) to 1230 and KI-GNN improving from 2915 (GNN) to 1379, outperforming both the existing methods (RF and GNN) and the expert-curated knowledge-only methods (C-RF: 4618, C-GNN: 2141). The KI-RF model showed a statistically significant improvement over both RF (p-value: 3.42e-03) and C-RF (p-value: 7.61e-16). Similarly, KI-GNN achieved significant improvements over both GNN (p-value: 4.69e-04) and C-GNN (p-value: 1.61e-08). These results demonstrate that combining experimental data with expert-curated knowledge yields better performance than using either data source alone. For AD DEE, KI-RF maintained a comparable median rank to RF (710 vs 718), while KI-GNN improved from 1306.5 to 1085. For AR DEE, KI-RF improved from 1779 to 1643, and KI-GNN improved from 4748 to 1642.

Table 1b reports median rank results for four other neurological diseases. For Microcephaly, KI-RF improved from 3688 (RF) to 2691, while KI-GNN showed a smaller improvement from 4748 (GNN) to 3898.5. In Intellectual Disability, KI-RF changed slightly from 2872 to 2896, while KI-GNN worsened from 3016 to 3771. Motor Neuron Disease presented the largest improvements, with KI-RF improving from 5305 to 669 and KI-GNN from 8551 to 407. For Cerebral Palsy, both models benefited significantly (KI-RF: from 2050 to 1450.5; KI-GNN: from 3338 to 1040).

Table 1c reports results for two non-neurological diseases to test generalisability. For Bleeding and Platelet Disorder, KI-RF achieved a median rank of 246, comparable to RF (249.5), while KI-GNN improved substantially from 6988 (GNN) to 104. For Arthrogryposis, KI-RF improved from 3488.5 to 2492.5 and KI-GNN from 6299 to 1828.5. The knowledge-only methods C-RF and C-GNN show varied performance across these diseases, further confirming that the integration of both data sources provides the most consistent improvements.

Overall, KI-RF and KI-GNN outperform both the existing methods and the knowledge-only methods in most cases, demonstrating the benefit of combining both data sources through the KIML framework. Possible causes for the worsened results are elaborated in the Discussion section.

#### 2.2.2 Biological Evaluation

All biological evaluations were conducted on the top 100-ranked predicted genes obtained by each method, representing the most confident predictions and a feasible number for downstream biological and clinical assessment. Only the evaluation results for DEE, the primary disease of focus, are presented in this section. The known DEE gene list was obtained from Genes4Epilepsy [17], using the version published in March 2025, to ensure that the predictions reflect the up-to-date information. AD and AR DEE results are provided in Supplementary Fig. 1-8, other results are in Supplementary Data 5 and 6.

##### Loss-of-function intolerance analysis using pLI scores

Loss-of-function (LoF) analysis was performed by evaluating the distribution of pLI (probability of loss-of-function intolerance) scores across candidate genes [18]. Disease genes tend to show higher pLI scores compared with the average protein-coding gene. We therefore expect disease genes discovered in the future to have a similar pLI distribution to known DEE genes; consequently, methods with better performance should yield predictions that more closely match the known DEE pLI distribution.

As shown in Fig. 6, the distribution obtained from KI-GNN is the most similar to the distribution of the known DEE genes, yielding the largest p-value (6.08e-01), followed by KI-RF (p-value: 4.05e-02). Comparing across the three categories, the existing methods RF and GNN, both show relatively narrow distributions concentrated in the upper range of pLI scores. The expert-curated knowledge-only methods C-RF and C-GNN, produce distributions with p-values of 9.0e-05 and 6.94e-03, respectively. The proposed Knowledge Inclusive methods achieve the closest alignment with the reference distribution: KI-RF demonstrates a broader distribution that shifts toward lower pLI values compared to both RF and C-RF, while KI-GNN shows a marked improvement over both GNN and C-GNN in matching the known DEE pattern. These results suggest that neither experimental data nor expert-curated knowledge alone is sufficient to fully recapitulate the pLI characteristics of known DEE genes, but their integration through the KIML framework yields distributions most consistent with the reference.

**Fig. 6:**
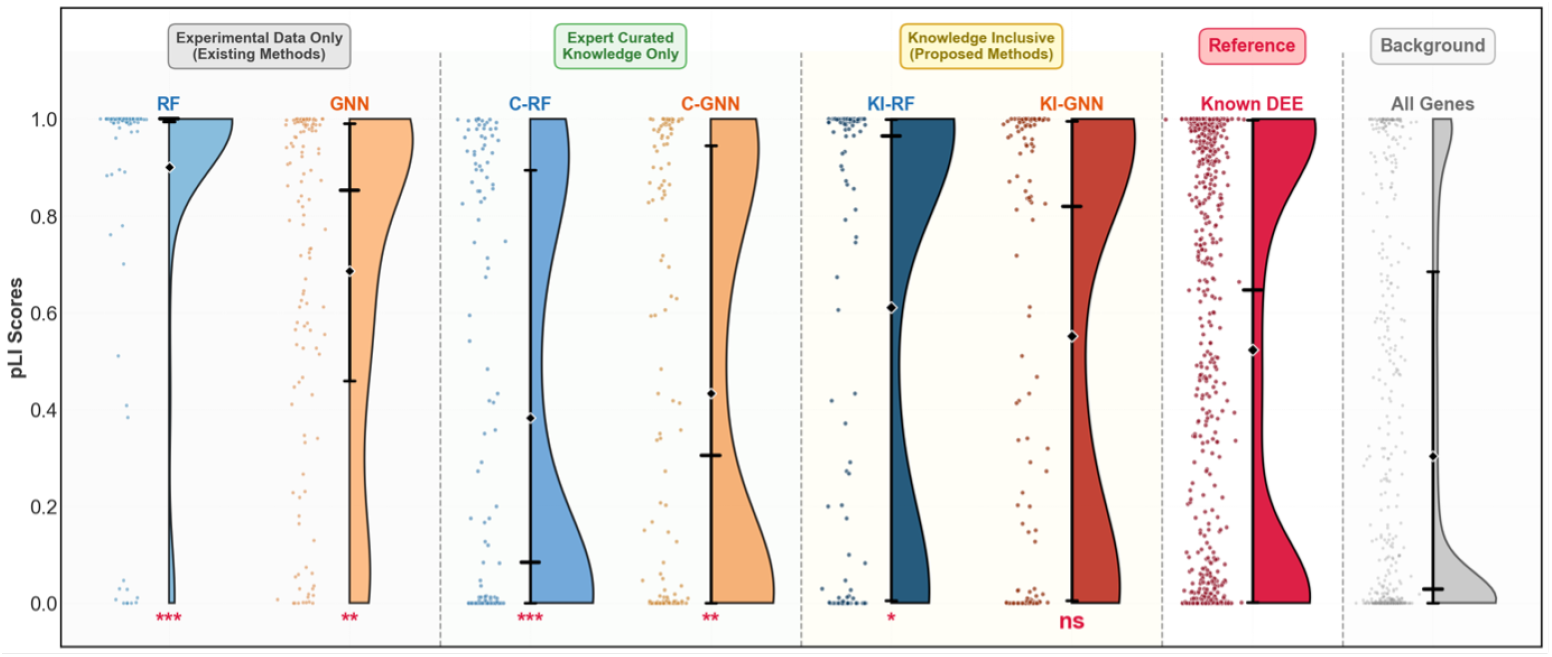
Distribution of probabilities of LoF intolerance (pLI) scores for candidate genes predicted by different methods, along with the distribution of known DEE genes (reference, in red) and all human genes (background, in grey). Existing methods (RF, GNN) are shown on the left, expert-curated knowledge only methods (C-RF, C-GNN) in the middle, and proposed Knowledge Inclusive methods (KI-RF, KI-GNN) on the right. The distributions were compared against known DEE genes using the Mann-Whitney U test, with significance levels indicated below each method (significance levels: *** *p <* 0.001, ** *p <* 0.01, * *p <* 0.05, ns = not significant).

##### Literature-based evaluation using phenotype–gene associations

To further evaluate the biological relevance of our predicted candidate genes, we performed a literature-based evaluation. The aim was to assess whether the prioritised candidate genes have been reported in the literature as associated with phenotype characteristics of DEE. If our predicted candidate genes are truly relevant to DEE, we would expect to find evidence in the literature linking them to these phenotypes. For DEE, we used the phenotype terms “Seizure” and “Intellectual disability” from the Human Phenotype Ontology (HPO), with identifiers HP:0001250 and HP:0001249, which correspond to two commonly observed clinical features [19].

To systematically search for such evidence from the literature, we used AMELIE [20], a phenotype–gene association search tool as previously employed by [12]. AMELIE compares genes against user-specified phenotypes and produces a score ranging from 0 to 100 for each gene, reflecting the strength of its association with the predefined phenotype terms (for DEE, i.e., HP:0001250 and HP:0001249) in the published literature.

Fig. 7 shows the violin plot of scores for the top 100-ranked candidate genes. The Mann-Whitney U test was used to statistically compare performance. Within the RF family, KI-RF achieves a significantly higher score distribution compared to RF (p-value: 3.04e-06), and similarly, KI-GNN significantly outperforms GNN (p-value: 4.56e-09), confirming the benefit of incorporating expert-curated knowledge. The comparisons between KI-RF and C-RF (p-value: 8.45e-02) and between KIGNN and C-GNN (p-value: 3.24e-01) do not reach statistical significance, indicating that expert-curated knowledge is the primary driver of capturing disease phenotype-gene associations. This highlights the advantage of incorporating expert-curated knowledge, which captures phenotype–gene associations less effectively represented in experimental data.

**Fig. 7:**
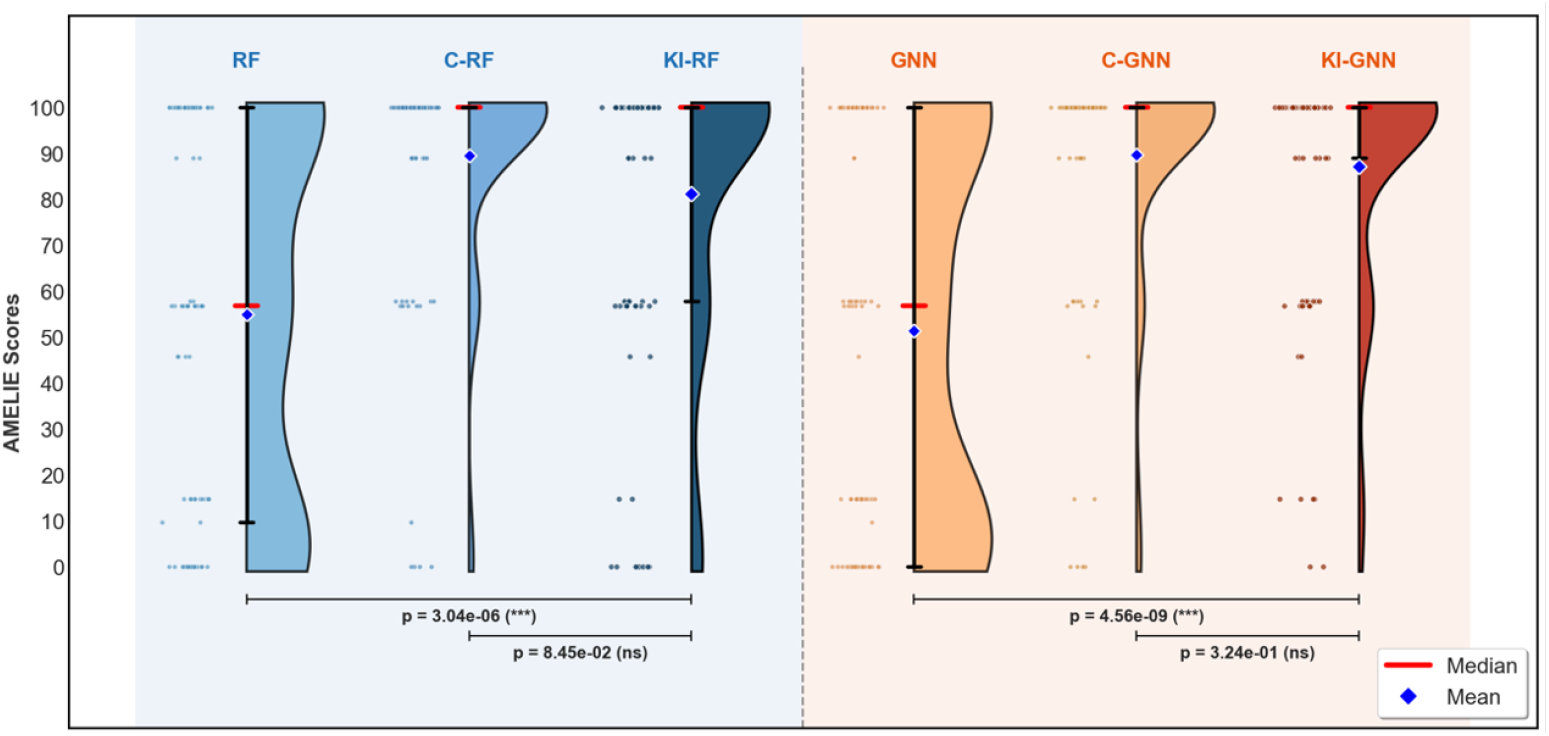
AMELIE scores for the top 100-ranked candidate genes across different methods. The red line indicates the median score, and the blue diamond marker indicates the mean score. Pairwise comparisons were performed using the Mann-Whitney U test, with corresponding p-values shown above the plot (significance levels: *** *p <* 0.001, ** *p <* 0.01, * *p <* 0.05, ns = not significant).

##### Phenotype enrichment analysis using Human Phenotype Ontology

Phenotype enrichment was performed on the top 100-ranked candidate genes derived from each method, using all protein-coding human genes as the reference list. The standardised vocabulary of phenotypic abnormalities was obtained from the Human Phenotype Ontology (HPO) [13].

Fig. 8 shows the top 10 enriched phenotypic terms identified by each method. The existing methods using only experimental data show no significantly enriched terms (all FDR *>* 0.05). In contrast, methods incorporating expert-curated knowledge, including both the Knowledge-only (C-RF, C-GNN) and Knowledge Inclusive (KI-RF, KI-GNN) methods, identify significantly enriched DEE-related phenotypes such as “Neurodevelopmental abnormality”, “Abnormality of mental function” and “Abnormal central motor function”. Both categories achieve significance (FDR *<* 0.05) across their top enriched terms, confirming that expert-curated knowledge enables the identification of core DEE phenotypic associations.

**Fig. 8:**
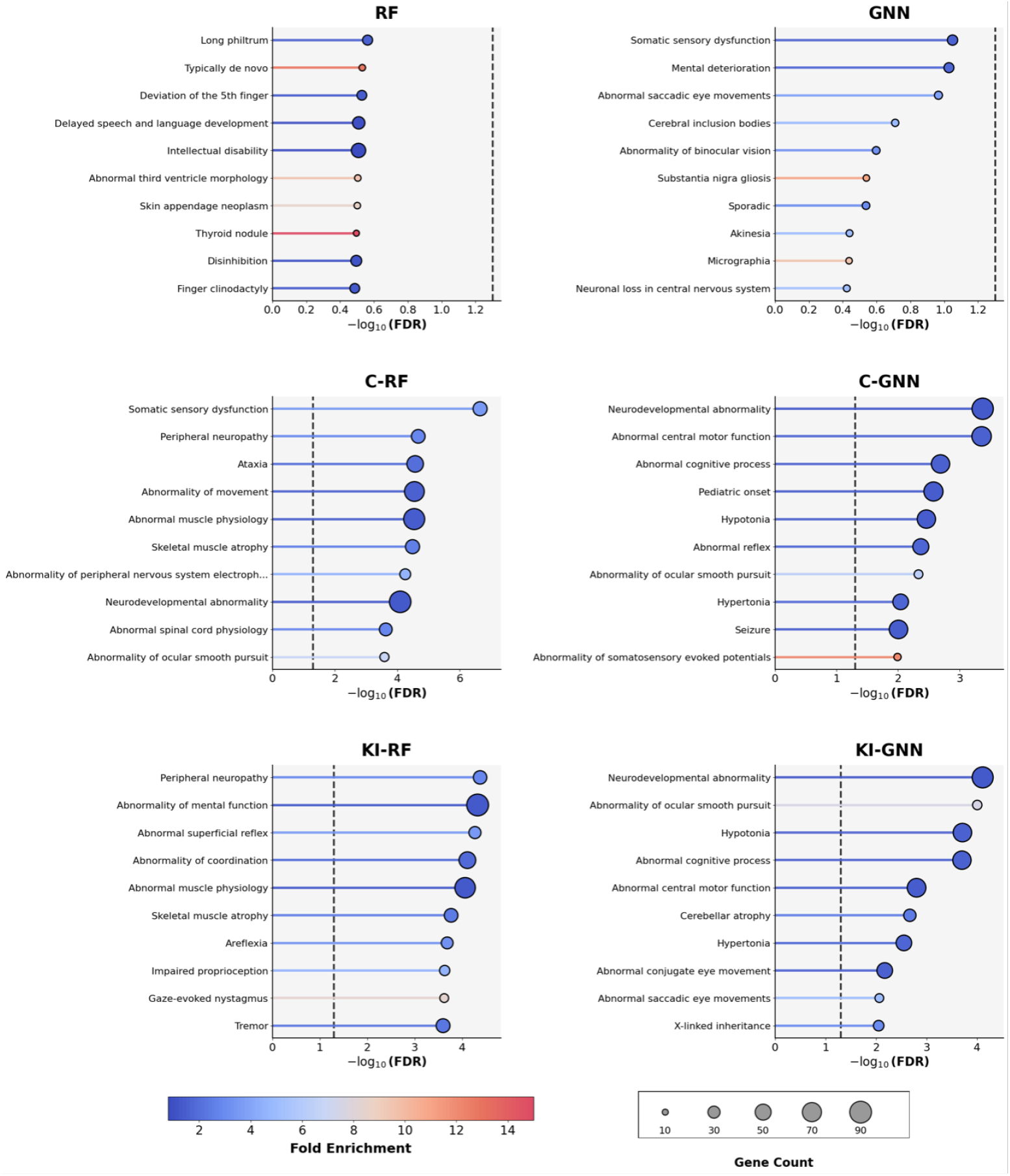
Phenotype enrichment evaluation results. The top 10 enriched phenotype terms from each method are shown as horizontal bar plots. Bar lengths represent the statistical significance of enrichment (− log(FDR)), with the vertical dashed line indicating the FDR cutoff (FDR = 0.05; −log(FDR) = 1.3). Circle sizes at the bar ends denote the number of candidate genes (out of 100) associated with each term, while colours indicate the fold enrichment values.

##### Gene Ontology enrichment analysis on biological processes

Gene Ontology (GO) enrichment was performed using the PANTHER classification system [21] via the Gene Ontology website with all protein-coding human genes as the reference list. The analysis focused on the biological process aspect, which provides the most direct insight into disease-related biological functions.

Fig. 9 presents the top 10 enriched GO biological process terms identified by each method. RF and GNN both capture some neurological terms, such as “Synaptic signalling” and “Regulation of trans-synaptic signalling”, but their enrichment profiles also include broader cellular processes such as “Cellular component organisation” and “Regulation of biological quality”. The knowledge-only methods C-RF and C-GNN similarly identify “Nervous system development” and “Synapse organisation”, alongside more generic metabolic and cellular terms. The Knowledge Inclusive methods KI-RF and KI-GNN show the highest concentration of neurodevelopment-related terms among their top enrichments: KI-RF identifies “Nervous system development”, “Neuron development” and “Generation of neurons”, while KI-GNN captures “Synapse organisation”, “Nervous system development” and “Regulation of membrane potential”, processes closely associated with neurodevelopmental and epileptic disorders.

**Fig. 9:**
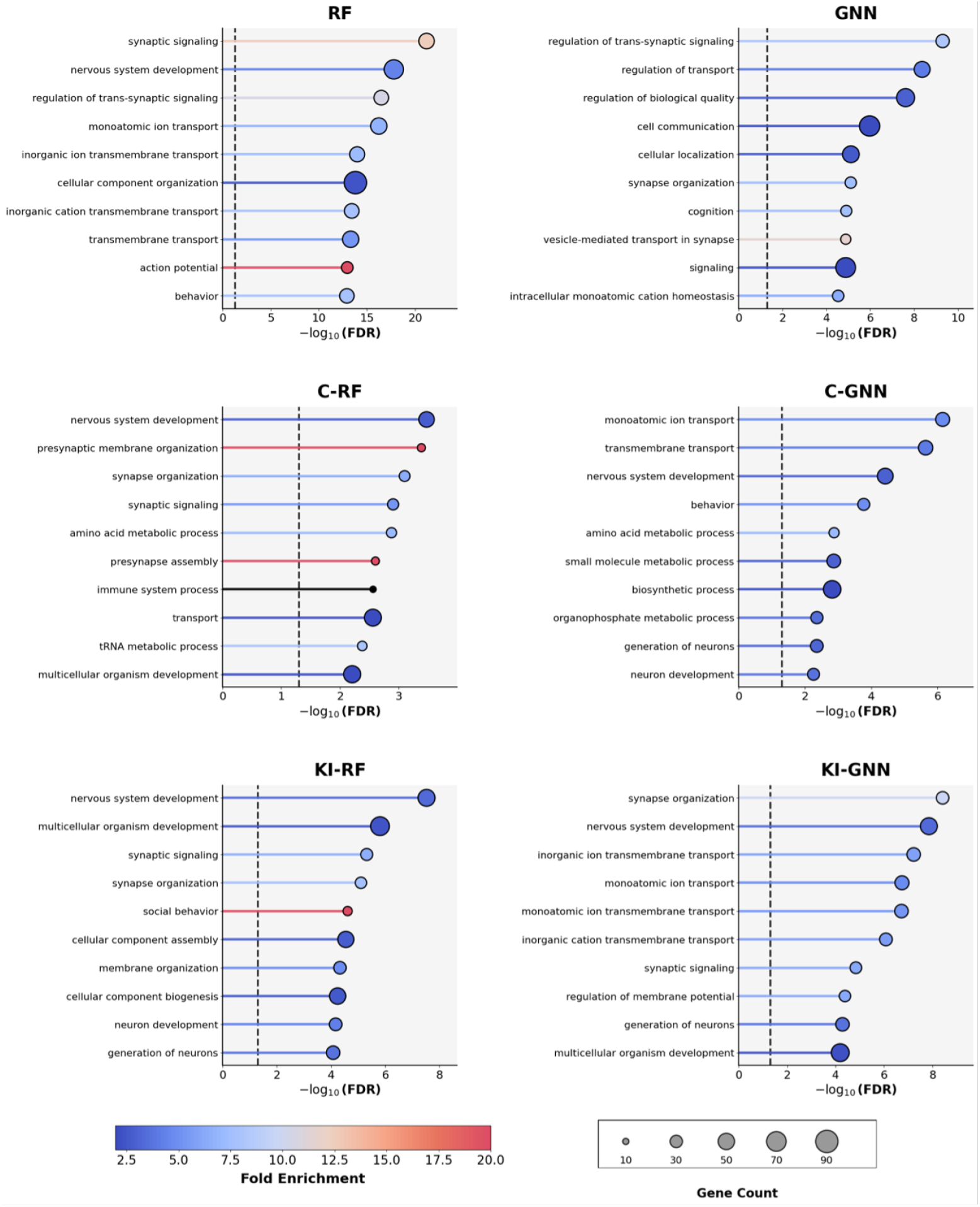
Gene Ontology enrichment evaluation results. The top 10 enriched Gene Ontology (GO) terms from each method are shown as horizontal bar plots. Bar lengths represent the statistical significance of enrichment (− log(FDR)), with the vertical dashed line indicating the FDR cutoff (FDR = 0.05; − log(FDR) = 1.3). Circle sizes at the bar ends denote the number of candidate genes (out of 100) associated with each GO term, while colours indicate the fold enrichment values.

### 2.3 Example of Model Interpretation based on KI-GNN

It is important to understand which biological features are most relevant to gene prioritisation results. This section, therefore, presents an example of model interpretation using the KI-GNN model. Fig. 10a highlights the landscape around the query gene *TSC2*, which we selected for detailed analysis because it is predicted as a high-confidence DEE candidate gene. Notably, *TSC2* ranks 10th in KI-GNN, while it is only ranked 1153rd in the GNN model, illustrating that the incorporation of expert knowledge improves gene prioritisation performance for this case. There are only 139 unique genes connected to TSC2 via edges with importance scores greater than 0.65. In Fig. 10a, the surrounding nodes represent the top 20 genes that contribute most to the prediction outcome. Several of the directly connected genes (red nodes) are already identified as DEE genes, such as *PACS1, UBA5* and *TPK1*, highlighting the biological relevance of this neighbourhood. Furthermore, the diversity of edge types indicates that *TSC2* is involved in multiple biological pathways, including “Signalling by Insulin receptor”, “Signalling pathways” and “BDNF signalling pathway”, reflecting its cross-functional role in neuronal processes.

**Fig. 10:**
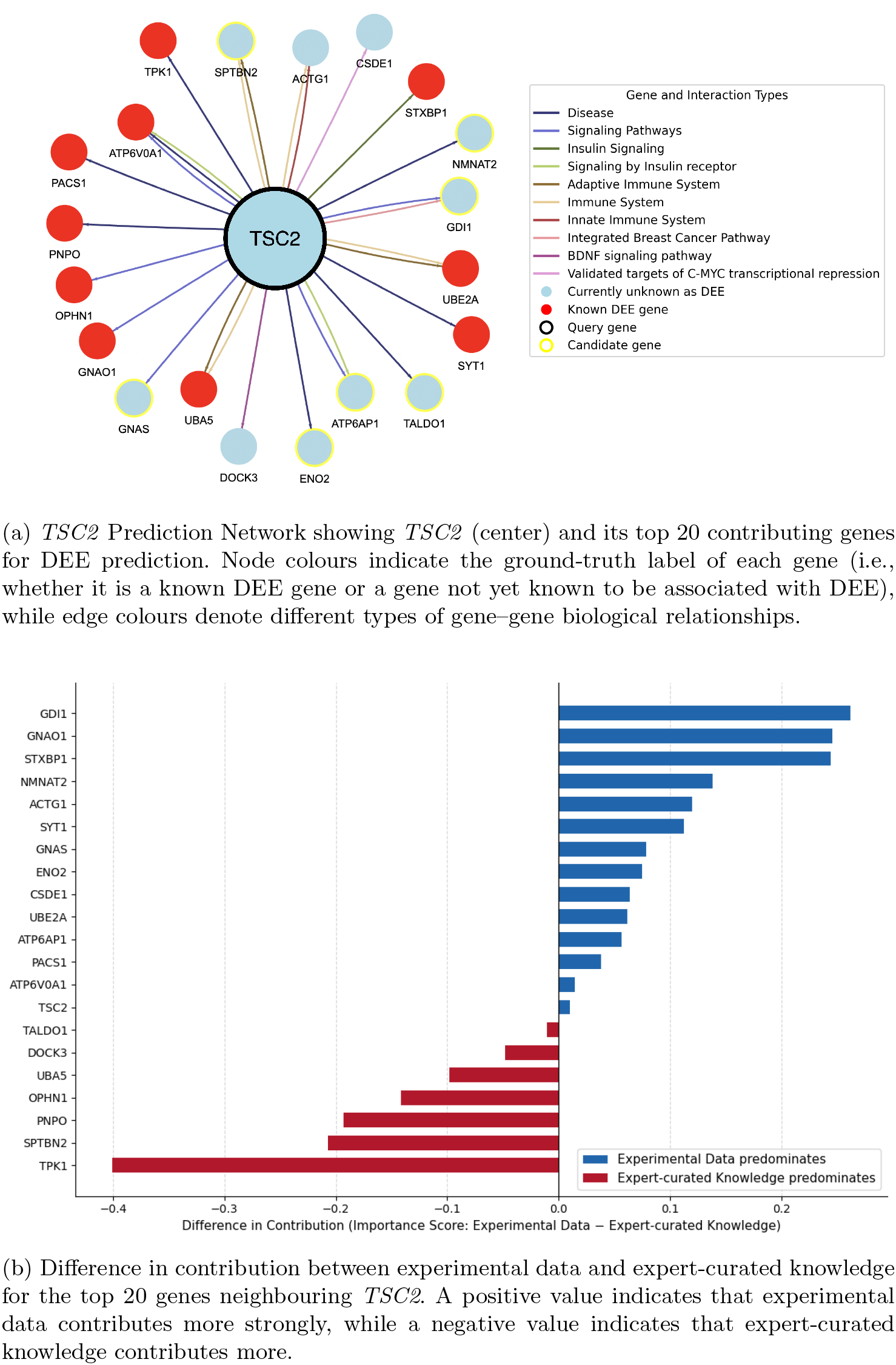
Model interpretation examples for the query gene *TSC2*.

**Fig. 11:**
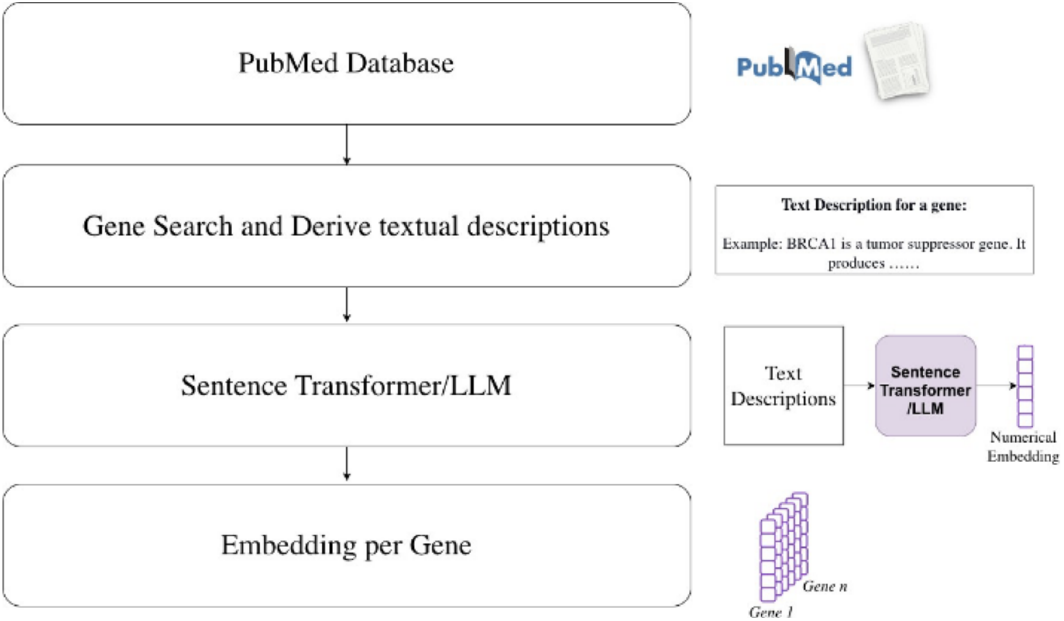
Pubmed Embeddings Generation

Fig. 10b further illustrates that among these top 20 contributing genes, experimental data and expert-curated knowledge contribute differently across individual genes. Genes such as *GDI1, STXBP1* and *GNAO1* are predominantly characterised by experimental data, suggesting their relevance to *TSC2* is largely driven by shared gene expression patterns. In contrast, genes such as *TPK1, SPTBN2* and *PNPO* are more strongly supported by expert-curated knowledge, indicating their association with *TSC2* is grounded in curated biological evidence. This demonstrates that the model draws on multiple sources of evidence across neighbouring genes, rather than relying uniformly on a single data type, thereby underscoring the value of knowledge integration in producing biologically interpretable predictions.

## 3 Discussion

In this paper, we propose a knowledge-inclusive machine learning pipeline that combines expert-curated knowledge with experimental data to prioritise candidate disease genes, while also providing interpretable explanations for its outputs. We demonstrated that the performance of disease gene prioritisation is significantly improved by including expert-curated knowledge as input to ML methods, which typically only use experimental data, as evidenced by both biological and temporal evaluations on DEE and six other diseases.

Even though we highlight the importance of expert-curated knowledge, it is important to note that using expert-curated knowledge alone did not give optimal results in our experiments. This can be attributed to the distinct biases inherent in each data source. Expert-curated knowledge is derived from published literature and established databases and is therefore biased toward well-studied genes and known disease associations, with limited coverage of novel or less-characterised genes. Experimental data, on the other hand, capture gene-level measurements without prior assumptions about disease relevance, but may lack the disease-specific context needed to distinguish true disease genes from broadly expressed or functionally active genes. These biases explain why neither data source alone is sufficient: expert-curated knowledge provides phenotype-gene association context that experimental data lacks, while experimental data offers unbiased coverage that compensates for the knowledge gaps in curated sources. The Knowledge Inclusive framework leverages this complementarity, allowing the models to ground novel discoveries from experimental data within the established biological context provided by expert-curated knowledge.

An interesting observation was made in inheritance pattern-specific DEE disease gene prioritisation. AD DEE gene prediction already had strong performance even without the inclusion of expert-curated knowledge, and the inclusion of expert-curated knowledge yielded only marginal improvements (Table 1). In contrast, AR DEE gene predictions benefited more from the inclusion of expert-curated knowledge. This suggests that characteristics captured in the experimental data, such as gene expression profiles, may provide stronger predictive signals for AD disease genes, whereas AR disease genes are more effectively prioritised by contextual knowledge that encodes mechanistic insights not directly reflected in experimental measurements.

GNNs require graph-structured data that accurately represents the specific biological context of disease gene prioritisation. To meet this requirement, we implemented a customisation pipeline of our framework (Supplementary Fig.9), where we apply a greedy method to select the optimal combination of datasets for DEE (since our code is publicly available, it can also be applied to additional diseases to customise the data and model selection process). Knowledge graphs provide a natural companion to our knowledge-inclusive pipeline in this regard due to being curated from scientific knowledge and innately containing a GNN-friendly format, i.e., edge data.

The PubMed literature-derived gene descriptions presented in this paper were processed to obtain text embeddings using a sentence transformer [10]. To avoid information leakage in the temporal-split evaluation (where the ML model was trained on disease genes available up to 30th Sep 2022 and evaluated on its ability to predict disease genes discovered after this partition date), both the literature corpus and the pre-trained sentence transformer were restricted to sources published before 2022. We also evaluated the ChatGPT embedding model [22], released in Dec 2022, with full results and a comparison against the sentence transformer provided in Supplementary Table 5. Overall, the ChatGPT embeddings outperformed the sentence transformer. However, since the release date of the ChatGPT embedding model (Dec 2022) overlaps by three months with the evaluation window (Sep 2022 to the present), we present only the sentence transformer results in the main text, which still demonstrate strong performance. It should be noted that this release date restriction is only for the purpose of temporal-split evaluation to avoid information leakage. In real-world applications, disease gene prioritisation models will be trained using the most up-to-date experimental data and expert-curated knowledge, together with all disease genes discovered to date. They do not need to employ temporal partitioning when prioritising candidates for future discovery, thereby mitigating the practical impact of this restriction.

As shown by the results of the temporal-split evaluation, our proposed KIML methods (KI-RF and KI-GNN) achieve better performance in most disease scenarios. However, we observed poorer performance for Intellectual Disability. This observation is likely influenced by two factors. First, a single global partition date (30th September 2022) was applied across all diseases as the sentence transformer model was restricted to literature published before 2022 to avoid information leakage. However, since different diseases have distinct discovery timelines, the number of genes identified before and after the partition date varies. If the number of disease genes identified before the partition date is very low, the model will underperform due to the lack of positive examples for training. Further, the ratio of known disease genes between the training and validation sets can become either too large or too small for certain diseases, affecting the stability and comparability of model performance (the detailed train/validation set sizes are provided in Supplementary Table 2). This imbalance affects the distribution of ranks given to disease genes in the validation set and may bias performance evaluation. Despite this limitation, the temporal-split evaluation primarily serves to demonstrate that the machine learning models can learn effectively from past experimental data and expert-curated knowledge. Second, this observation may also be related to the data integration strategy used in our framework. As illustrated in Fig. 3, these two types of information are concatenated directly, implicitly assuming equal importance across all diseases. However, the relative contribution of experimental data and expert-curated knowledge is likely to be disease-dependent. For example, for some diseases, experimental data may be noisy or less informative, while expert-curated knowledge may provide more reliable and discriminative information (e.g., the median rank results of GNN-based methods for Motor Neuron Disease in Table 1). In such cases, assigning higher weights to expert-curated knowledge could lead to improved prioritisation performance. Therefore, incorporating disease-specific weighting or adaptive fusion strategies represents a promising direction for future work to further enhance the flexibility and robustness of the KIML framework.

The biological evaluation results demonstrate that with the inclusion of expert-curated knowledge, the genes predicted by the models more closely resembled the characteristics of known disease genes, in terms of their loss-of-function intolerance scores, phenotype, and gene ontology. Although the expert-curated knowledge derives from the literature, which already contains extensive knowledge about these genes and could raise concerns about potential information leakage, it is important to note that we did not explicitly direct the machine learning model to focus on any particular attributes or characteristics. Instead, the model itself was able to identify and learn the patterns that distinguish disease genes, indicating that it has captured the useful underlying features.

To further improve the performance of gene prioritisation, future work will focus on refining the construction of edges in GNNs. In the current work, the edge connections are derived from an expert-curated knowledge graph, HetioNet [7], which integrates information from 29 heterogeneous datasets covering diverse biological aspects such as PPIs, pathways, and phenotypic associations. Although this provides a comprehensive foundation, the knowledge captured is limited by the scope of the underlying datasets. To address this limitation, future work can incorporate PubMed literature-based edge connections. Specifically, large language models can be employed to extract information from biomedical literature, enabling the generation of edges that reflect the comprehensive scientific discoveries. In addition, models can be further improved by integrating additional experimental data, such as DNA methylation data. Since biological systems are inherently complex, incorporating epigenetic information beyond gene-level features can provide additional insights. By adding additional layers of information, the model is more likely to capture more biologically relevant patterns. Such integration is expected to enhance the biological interpretability of the KIML pipeline while also improving prediction performance. Furthermore, our knowledge-inclusive approach has the potential to accommodate other sources of expert-curated knowledge, such as mathematical models and equations within the domain of gene studies. These mathematical equations can be incorporated as constraints or features within the ML models, guiding the learning process.

Regarding the interpretability of the GNN and KI-GNN models in the pipeline, our explainability approach for understanding model decisions behind disease gene prioritisation employs the GNNExplainer [23]. We provide explanations for the model’s top-ranked candidate genes, with each explanation tracing the important pathway links that support the gene’s classification as disease-causing. One limitation inherent in current graph explainability methods, is an effect known as the introduced evidence problem [24]. These methods test the importance of connections by partially altering them, which can unintentionally modify the underlying data by creating new patterns that were not originally present. We plan to improve our explanations for disease gene prioritisation by incorporating techniques such as graph transformer networks [25] and reinforcement learning [26]. These methods generate explanations by identifying specific connections and pathways that already exist in the biological network, rather than modifying the network itself, which avoids the introduced evidence problem and ensures the explanations remain faithful to the actual biological relationships.

## 4 Methods

### 4.1 Information-Inclusive Machine Learning Framework

We propose the concept of expert-curated knowledge inclusive Machine Learning (InclusiveML). Within a biomedical context, such real-world data used in Machine Learning typically corresponds to experimental data collected from biomedical samples, conditions, or cohorts. Expert-curated knowledge refers to any background knowledge related to experimental data captured as text, mathematical equations, specific data preprocessing methods, knowledge graphs, multimedia, etc. Expert-curated knowledge ensures the provision of all available knowledge about data which may include theoretical foundations/reality for the Machine Learning model to build upon. In our InclusiveML pipeline for disease gene prioritisation, we draw primarily on two types of expert-curated knowledge, which we add in addition to multiple types of experimental data sources.

#### 4.1.1 Expert-Curated Knowledge

Published scientific literature (expert-curated by the scientific review process) provides valuable context for individual genes (node data) regarding previous experimental and biological findings. An expert-curated Knowledge Graph provides context for inter-gene interactions (edge data), allowing the methods that can handle edge data to be tested.

##### PubMed literature-derived gene descriptions

The initial step in deriving gene descriptions from scientific literature in the PubMed database involved identifying publications that reference specific genes. We iterated through all the PubMed publications and searched for references to genes in the abstract section and in the title. In this process, we considered multiple naming conventions of genes, such as gene symbols (e.g., SCN1A, BRCA1), Ensembl IDs (e.g., ENSG00000144285, ENSG00000012048), NCBI Gene IDs (e.g., 6323, 672), and HGNC IDs (e.g., HGNC:10585, HGNC:1100). This step generates a list of papers for each gene. We filtered out the publications published after the temporal partition date (September 2022). We then retrieved the corresponding titles and abstracts for each gene via the PubMed API. The concatenated titles and abstracts from associated papers per gene were considered expert-curated knowledge or descriptions for individual genes *L*_*i*_ for gene *g*_*i*_. These gene descriptions were then converted into numerical embeddings using a pretrained language transformer model. This model can capture the semantic meaning of text in a compact vector format:

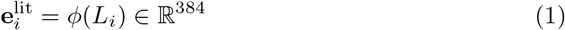

where *ϕ* is the sentence-transformer model and 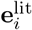 is the literature embedding for gene *g*_*i*_. Two alternative architectures were evaluated for this purpose: the sentence-transformer “all-MiniLM-L6-v2”[27], which provides a vector of 384 dimensions, and the OpenAI text embedding model “text-embedding-ada-002”[28], which provides a vector of 1536 dimensions. The sentence-transformer “all-MiniLM-L6-v2” was trained in July 2021, well before the partition date in our temporal-splite evaluation method.

This guarantees that there is no information leakage present, i.e., an overlap between expert-curated training data and experimental test data of the sentence transformer. The OpenAI text embedding model “text-embedding-ada-002” was published in December 2022, which is 3 months after our partition date in the temporal-split evaluation, and may lead to information leakage. Therefore, we only present the results obtained from sentence-transformer “all-MiniLM-L6-v2” embeddings in this paper. However, a comparison between the results obtained from these two different language embedding models is in Supplementary Table 5.

##### Knowledge graph

A knowledge graph is a conceptual representation of a group of entities (nodes) and their relationships (edges) in the form of a graph 𝒢 = (𝒱,ℰ) where represents the set of vertices (nodes) and ℰ represents the set of edges. Knowledge graphs are formulated by collecting and curating information from multiple data sources. In this study, we utilise HetioNet [7], a biomedical knowledge graph that encapsulates relationships uncovered by thousands of studies over the last half-century. It is important to note that HetioNet was published in 2017, before the partition date, making it appropriate to integrate without filtering for our temporal-split evaluation protocol. Hetionet contains multiple types of nodes, including Gene, Disease, Biological Process, Pathway, etc. These multi-typed nodes are connected based on their relationships. For example, if a particular gene is participating in a biological pathway, there is an edge between the respective gene and the pathway. In our study, we derived a graph of the gene-gene relationship using this knowledge graph. We constructed 𝒱 = *G* where *G* = {*g*_1_, *g*_2_, …, *g*_*N*_} is the set of all protein-coding human genes with *N* ≈ 20,000. For edges (*g*_*i*_, *g*_*j*_, *σ*) ∈ ℰ, if two gene nodes are connected to the same pathway node, we add an edge between those two gene nodes in the graph we derive with relation type *σ* ∈ Σ where Σ is the set of relation types. Similarly, we derived different gene-gene graphs using biological processes, diseases, molecular functions, etc. In our study, we used these derived graphs to train GNN-based models.

Our ablation study on the effectiveness of each of these derived graphs in DEE gene prioritisation is in Supplemantary Table 6.

#### 4.1.2 Experimental Data

##### scRNA-seq expression data

Single-cell RNA-seq expression data was downloaded from human fetal cortical samples as described in a previous publication [29] and used in Mantis-ml [12]. The dataset includes four samples collected across an 8-week period during mid-gestation, comprising 57,868 single-cell transcriptomes. Precomputed average gene expression profiles for major cell types across four developmental time points, as provided in Mantis-ml, were directly used in this study to construct features 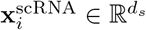 where *d*_*s*_ is the dimension of single-cell expression features.

##### Protein-Protein Interaction (PPI) dataset

The Protein-Protein Interaction (PPI) dataset we utilised is the Biological General Repository for Interaction Datasets (BioGRID) [30], which is an open-access database that houses genetic and protein interactions from primary biomedical literature for all major model organism species and humans. BioGRID contained 2.248 million non-redundant interactions derived from 87,206 publications.

##### Developmental transcriptomes in the brain

Developmental transcriptomes in the brain were obtained from BrainSpan (RNA-Seq Gencode v10 summarised to genes) [31]. This dataset consists of bulk RNA sequencing data in 26 brain sub-regions in 11 developmental stages, from embryonic stages to adulthood, with the sample size of each region ranging from 229 to 8,244 in total. The developmental stages were grouped into five periods: prenatal (4–37 post-conception weeks), infancy (birth–18 months), childhood (19 months–11 years), adolescence (12–19 years), and adulthood (20–60+ years), based on the criteria used by BrainSpan. Since DEE typically manifests at very early ages and affected individuals may not survive into adulthood, we restricted our analysis to the first four periods and excluded the adulthood stage, yielding features 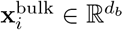 where *d*_*b*_ is the dimension of bulk RNA-seq features.

A co-expression network was then constructed using the developmental transcriptomes from the first four stages (excluding adulthood). The Spearman’s rank correlation coefficient *ρ*_*ij*_ was calculated to quantify gene-gene co-expression between genes *g*_*i*_ and *g*_*j*_. We tested several |*ρ*| thresholds (e.g., 0.5 and 0.8) and selected |*ρ*| = 0.8 as the threshold *τ*, as it resulted in the best performance in the subsequent analysis. Genes *g*_*i*_ and *g*_*j*_ are connected if |*ρ*_*ij*_| *> τ*.

#### 4.1.3 Labels

For DEE gene prioritisation, we used Genes4Epilepsy [17] as the source of known DEE gene labels (available at: github.com/bahlolab/genes4epilepsy). This curated database represents a set of actively curated monogenic epilepsy genes, which is updated twice a year. The first release (2022-09) contained 926 genes associated with epileptic phenotypes, including 825 genes specifically linked to DEE. In the 2025-03 version, this list had expanded to 1,040 genes, of which 937 were associated with DEEs. For other diseases, labels were taken from disease gene panels in PanelApp [32]. PanelApp is a crowdsourced knowledge base that publishes curated gene lists for many genetic diseases. In our work, the data partition date was set to 30th September 2022 (*t*_0_ = September 30, 2022). This date is chosen to provide a fair balance between the number of training and validation genes, particularly for our primary disease of focus, DEE. Further information about the training and validation set sizes for each disease is provided in Supplementary Table 2.

#### 4.1.4 Gene Prioritisation Models

##### Random Forest (RF)

We employed the model from Mantis[12], a random forest-based gene prioritisation framework. Originally, it uses general (e.g., GenomeAd, Genic intolerance scores), tissue-specific (e.g., Human phenotype ontology, GTEX genotype tissue expression), and disease-focused (e.g., chronic kidney disease, cardiovascular disease) experimental data sources to prioritise candidate disease genes. The Mantis framework handles the preprocessing of these datasets. To evaluate the effectiveness of expert-curated knowledge, we concatenated the PubMed embeddings 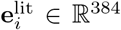 derived from PubMed literature, single-cell RNA-seq features 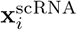, bulk RNA-seq features 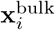 from BrainSpan developmental transcriptomes, and Mantis preprocessed features to form a unified feature vector. Specifically, the Mantis framework integrates genomic, functional, and disease-specific information from diverse public resources, including genetics databases (e.g., gnomAD, ExAC), genic intolerance scores (e.g., RVIS and MTR), functional annotations (e.g., GOA, MSigDB), gene expression and QTL datasets, GWAS resources, and disease-specific databases (e.g., CKDdb) to generate features 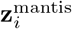. The concatenated feature vector is:

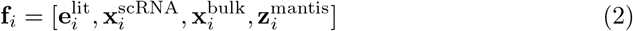

The Random Forest prediction is computed through ensemble averaging:

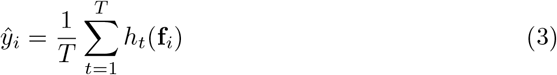

where *h*_*t*_ are individual decision trees and *T* is the number of trees in the ensemble.

##### Graph Neural Network (GNN)

We utilised the model from SPEOS [3], a positive-unlabelled (PU) graph representation learning framework for disease gene prioritisation. SPEOS models a heterogeneous graph where nodes represent all protein-coding human genes (*>* 20,000), with node features integrating experimental data (e.g., gene expression values) and expert-curated knowledge (e.g., PubMed-derived functional descriptions). Node representations are initialized by concatenating PubMed literature embeddings 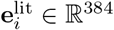 with single-cell RNA-seq features 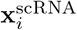:

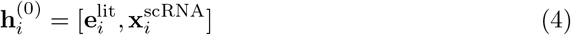

Edges capture diverse biological relationships, including those derived from experimental data (e.g., protein–protein interactions, coexpression networks) and curated knowledge graph edges (edges derived from expert-curated knowledge). Among the available base classifiers, we used FiLM [33], which demonstrated the best performance compared to other alternatives in SPEOS. In addition, the nested cross-validation ensemble strategy of SPEOS ensures robust and reliable performance.

The model employs message passing across *L*_max_ layers. At each layer *ℓ* ∈ {1, …, *L*_max_}, node representations are updated through aggregation and update functions. The aggregation function AGG^(*ℓ*)^ collects information from neighboring nodes:

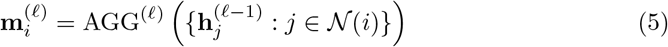

where 𝒩 (*i*) denotes the set of neighbors of gene *i*. The update function UPD ^(*ℓ*)^ combines the aggregated neighborhood information with the node’s own representation:

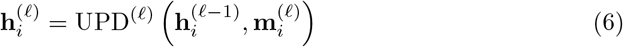

After *L*_max_ layers of message passing, final predictions are obtained via:

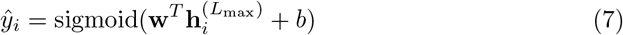

where **w** is a weight vector and *b* is a bias term.

The input datasets used for each method are summarised in Supplementary Table 1. RF and KI-RF share the same experimental data, as do GNN and KI-GNN, providing a fair basis for evaluating the performance improvements introduced by the KIML pipeline. The RF-based methods (RF and KI-RF) and the GNN-based methods (GNN and KI-GNN) were evaluated on the experimental datasets used in Mantis and SPEOS, respectively, allowing a direct quantification of the improvements over these existing methods. We also evaluated each method on alternative experimental datasets (Supplementary Table 3), which resulted in suboptimal results compared to the original experimental datasets used by the respective methods.

#### 4.1.5 Dataset Customisation

The dataset customisation was conducted to identify the optimal combination of datasets that yields the best performance for disease gene prioritisation. A greedy strategy was applied: in the first round, all candidate datasets were included, and performance was evaluated using the temporal-split evaluation procedure, with median rank as the evaluation metric. In each subsequent step, one dataset was removed at a time, and the resulting performance was compared. The configuration with the best median rank was retained as the starting point for the next round. This iterative process continued until no further improvement could be achieved by excluding additional datasets. The final combination of datasets with the best median rank was selected as the optimal dataset set. The dataset customisation results for DEE using the GNN-based method are presented in Supplementary Data 2. This method of dataset customisation can be applied to any other prediction, including the prediction of genes implicated in other diseases.

### 4.2 Temporal-Split Evaluation

Temporal-split evaluation of disease genes depends on the progression of disease gene discovery by experimental means over time. Let *G* = *{g*_1_, *g*_2_, …, *g*_*N*_ *}* represent the set of all protein-coding human genes where *N* ≈ 20,000, and let *y*_*i*_ ∈ *{*0, 1*}* denote the binary label indicating whether gene *g*_*i*_ is known to be associated with the disease of interest. In our method, the DEE disease gene list was obtained from the Genes4Epilepsy GitHub repository [17] for training and validation purposes. They have been updating their DEE gene list twice a year since September 2022. The DEE gene list corresponding to the September 2022 version was used to annotate the genes for the training process, and the newly added genes in later versions up to March 2025 were used to validate the proposed InclusiveML disease gene prioritisation pipeline. Formally, the temporal partition date was set to *t*_0_ = September 30, 2022, such that:

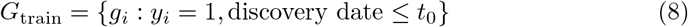

represents genes known before the partition date, and:

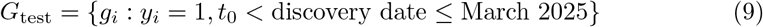

represents genes discovered after the partition date up to March 2025.

The other diseases used in our study were selected based on the availability of their disease gene lists on PanelApp [32]. In PanelApp, when new genes are discovered to be associated with a disease, they are added to the respective list of genes, accompanied by a timestamp. Using this timestamp, we derived the gene labels for the model training and validation protocol.

After the training process is completed, all genes not previously known to be associated with the disease are ranked in descending order based on their model-predicted scores:

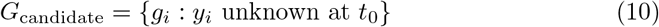

Then the ranks for the newly identified disease genes are obtained, and they are used to derive the mean and median ranks:

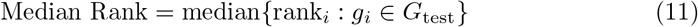

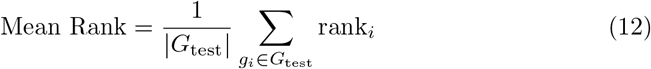

where rank_*i*_ ∈ *{*1, 2, …, |*G*_candidate_|} denotes the rank of gene *g*_*i*_ in the ordered list of candidates. Lower median and mean ranks indicate better performance. Further, the rank distribution of the newly identified disease genes across the ranked list is obtained to evaluate the statistical significance of improvements.

### 4.3 Biological Evaluation

All biological evaluations were conducted on the top 100-ranked predicted genes obtained by each method. These genes represent the most confident predictions and a feasible number for downstream biological and clinical assessment.

#### 4.3.1 Loss of Function Intolerance Analysis using pLI scores

The pLI score has values ranging from 0 to 1, where scores approaching 1 indicate intolerance to the variants. We compared the pLI score distributions between our predicted candidate gene sets (derived from different methods, such as the proposed KI-RF or KI-GNN) and a reference set of known DEE genes. The Kolmogorov–Smirnov test, a non-parametric method for comparing the overall distributions of two independent groups of samples, was used to statistically assess the similarity of these distributions across methods:

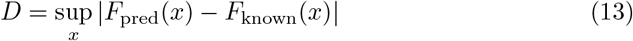

where *F*_pred_ and *F*_known_ are the cumulative distribution functions of predicted and known gene sets. This test was chosen because it makes no assumptions about the underlying distribution shape and is robust to outliers, making it particularly suitable for comparing pLI score distributions, which may not follow normal distributions. Methods yielding candidate gene sets with pLI distributions most similar to known disease genes were considered to demonstrate biological relevance and predictive accuracy for identifying disease genes.

#### 4.3.2 Enrichment Analysis

Enrichment analyses for phenotype and Gene Ontology (GO) were performed to test whether phenotypic features and biological processes are overrepresented in the top 100 candidate genes from each method, relative to their baseline frequency across all protein-coding human genes, using the hypergeometric test:

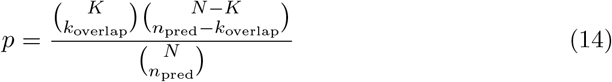

where *N* is the total number of genes, *K* is the number of genes with a given term, *n*_pred_ is the number of predicted candidates, and *k*_overlap_ is the overlap. For phenotype enrichment, we used the Human Phenotype Ontology (HPO), which provides over 13,000 standardized terms linking genes to phenotypic features, allowing systematic identification of enriched features. For GO enrichment, we used the GO knowledge base from geneontology.org. Statistical significance for both analyses was assessed with Fisher’s exact test and corrected for multiple testing using the Benjamini–Hochberg FDR.

#### 4.3.3 Literature-Based Evaluation

The literature-based evaluation was performed using AMELIE [20], a phenotype– gene association search tool that has also been applied in Mantis-ml [12]. AMELIE is designed to rank genes based on their relevance to a patient’s clinical phenotype by integrating information from scientific literature. The tool employs a machine learning classifier that assigns scores to gene-phenotype-article triples using 27 features across 6 categories: inheritance mode features, variant overlap features, phenotype similarity features, variant type features, article relevance features, and a priori pathogenicity features. A key component of the scoring mechanism is the phenotype similarity, which leverages Phrank, a tool designed to calculate similarity scores between patient phenotypes and disease descriptions in the literature using the Human Phenotype Ontology.

### 4.4 Model Interpretation

Explainable AI (XAI)[15] is used to make the gene prioritisation process transparent. Gene prioritisation methods must be grounded in scientific discovery and transparent validation to build clinical trust. Understanding why certain genes are highly ranked enables researchers and clinicians to make more informed decisions. Additionally, XAI supports model refinement by revealing why models succeed or fail, thereby guiding iterative improvement. Most of the existing literature on disease gene prioritisation does not consider explainability. Only a limited number of studies have tackled the need for XAI [3, 4], but the limitation is that their architectures either depend heavily on label propagation to refine explanations, which may reduce prioritisation accuracy or lack direct consideration of biological interaction details.

In our study, since the data used for disease gene prioritisation includes genegene interactions and since the SPEOS framework utilises Graph Neural Networks (GNN), we have used GNNExplainer [23] for explaining the model decisions. GNNEx-plainer uses a perturbation masking strategy to find the explanatory subgraph that best describes the reasoning behind the model’s prediction. For each prediction, GNNExplainer identifies a subgraph 𝒢_*S*_ = (𝒱_*S*_, ℰ_*S*_) that maximizes the mutual information(MI):

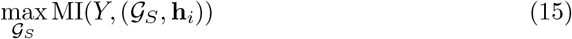

where *Y* is the prediction, 𝒱_*S*_ and ℰ_*S*_ are subsets of vertices and edges, and **h**_*i*_ is the node representation. This explanation includes importance scores for each interaction (edge importance), reflecting the strength of their influence on the prediction. In our study, we generated explanations for each candidate gene that appear in the top-ranked list to indicate the possible genes or interactions that influence the gene to be a candidate gene.

Since our training employs *K*-fold cross-validation, we aggregate edge importance across folds:

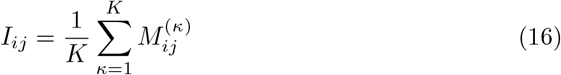

where 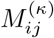 is the importance of edge (*i, j*) in fold *κ*. The Knowledge Graph-based gene-gene interactions[7] used in our method, such as pathway information, provide human-interpretability to the explanations generated by GNNExplainer.

#### 4.4.1 KIML DEE Gene Prioritisation Web Interface

The web interface created by us for the purpose of showcasing our model results for disease gene prioritisation in DEE is available freely. This tool displays the top-ranked disease genes discovered by the latest version of our model and the explanation obtained for each gene to be predicted as a disease-associated gene. Each explanation includes an interactive network highlighting the subnetwork responsible for the gene to be predicted as a disease gene. This interactive network has indications to show the known disease-causing genes, other candidate genes, and the gene-gene interaction types affecting the model’s decision, along with the interaction importance scores. Additionally, the explanation includes a description of the explanation graph and the same data in a tabular and downloadable format. Our web interface is hosted on the Streamlit platform, making it publicly accessible.

## Data Availability

All datasets used in this study are publicly available. DEE gene labels were obtained from Genes4Epilepsy (https://github.com/bahlolab/genes4epilepsy; 2022-09 release for training and 2025-03 release for validation). Disease gene panels for additional diseases were obtained from PanelApp (https://panelapp.genomicsengland.co.uk). Knowledge graph data were obtained from HetioNet (https://het.io, 2017 release). Protein-protein interaction data were obtained from BioGRID (https://thebiogrid.org). Bulk developmental transcriptomes were obtained from BrainSpan (https://www.brainspan.org, RNA-Seq Gencode v10). Single-cell RNA-seq data from human fetal cortical samples were obtained as previously described and used in Mantis-ml. Probability of loss-of-function intolerance (pLI) scores were obtained from the ExAC v0.3 constraint dataset (`forweb_cleaned_exac_r03_march16_z_data_pLI.txt`), available via the Broad Institute/gnomAD downloads portal. Phenotype terms were obtained from the Human Phenotype Ontology (https://hpo.jax.org). Gene Ontology terms were obtained from https://geneontology.org. PubMed literature was accessed via PubMed. No restrictions apply to data availability.

## Code Availability

KIML is open source, implemented in Python and available at: https://github.com/jayasankha1010/KIML.

## Author Contributions

Conceptualization: S.H., C.J.G., Y.X., S.S.; Model development and evaluation analyses: C.J.G., Y.X., R.R., S.S., M.B., M.F.B.; Method development: C.J.G., Y.X., R.R., S.S., D.S., T.M., A.H., S.H.; Implementation: C.J.G., Y.X., R.R., S.S.; Clinical and genetic expertise: M.C., T.J.O., S.P., S.F.B., I.E.S., J.G., M.B., M.F.B.; Paper writing: C.J.G., Y.X., R.R., S.S., A.H., S.H.

## Acknowledgements

We thank all the individuals for participating in this research. Funding for this project was provided by the NHMRC Synergy Grant (GA202712): “Integrative-omics” for precision medicine in the epilepsies.

