## supplementary figures and tables for "Knowledge Inclusive Machine Learning for Disease Gene Prioritisation"

**Supplementary Tables**

Supplementary Table 1. Number of disease genes in the training versus validation sets for Temporal-split validation (Partition date: September 30, 2022).

| Disease | Number of disease genes considered for training | Number of disease genes considered for validation |
| --- | --- | --- |
| DEE | 825 | 223 |
| Autosomal Dominant DEE | 291 | 73 |
| Autosomal Recessive DEE | 492 | 53 |
| Microcephaly | 267 | 46 |
| Intellectual Disability | 1923 | 221 |
| Motor Neuron Disease | 66 | 7 |
| Cerebral Palsy | 92 | 233 |
| Bleeding and Platelet Disorder | 115 | 15 |
| Arthrogryposis | 174 | 22 |

Supplementary Table 2. The input datasets used for each method.

| Types of data | RF | C-RF | KI-RF | GNN | C-GNN | KI-GNN |
| --- | --- | --- | --- | --- | --- | --- |
| Human Ontology | 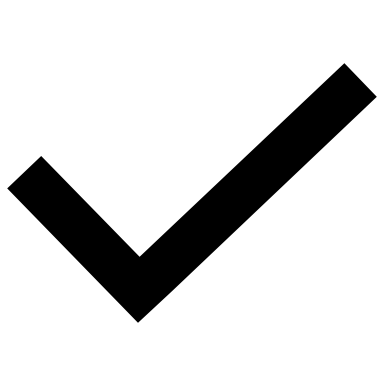 |  | 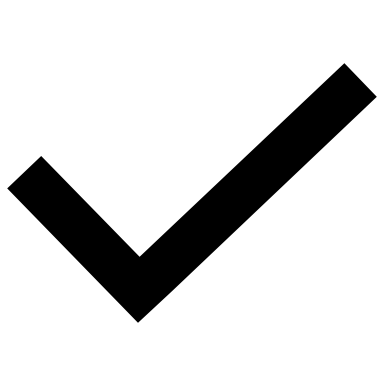 |  |  |  |
| GWAS | 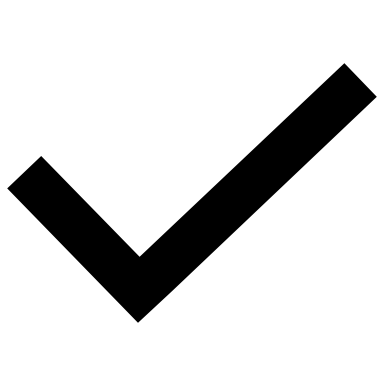 |  | 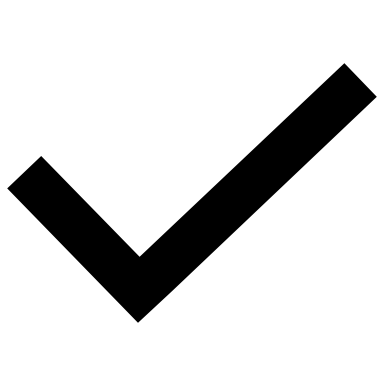 |  |  |  |
| GnomAD | 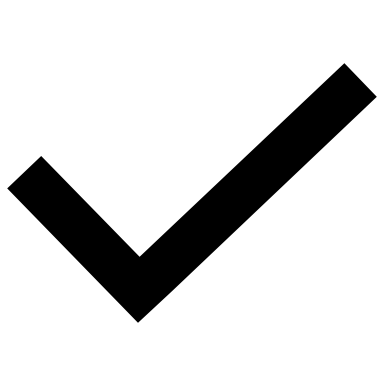 |  | 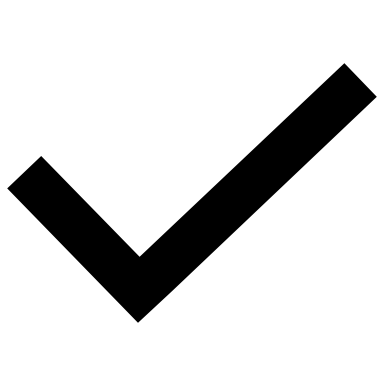 |  |  |  |
| ExAC | 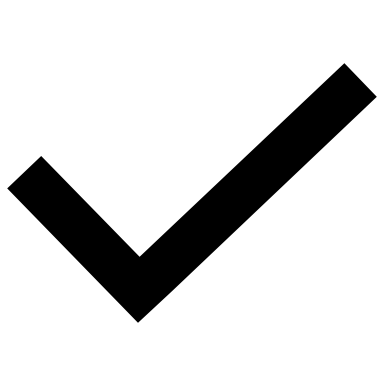 |  | 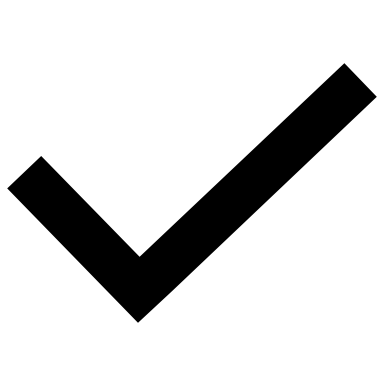 |  |  |  |
| Genic-intolerance scores  (RVIS, MTR) | 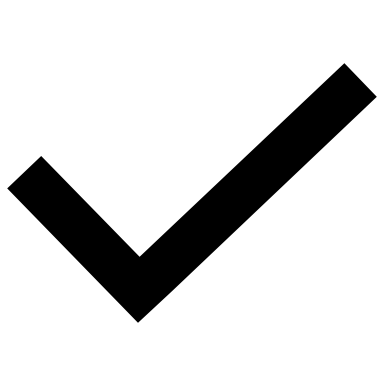 |  | 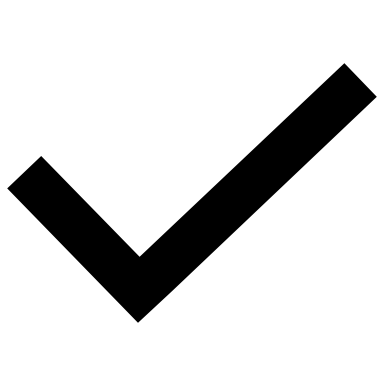 |  |  |  |
| InWeb_IM | 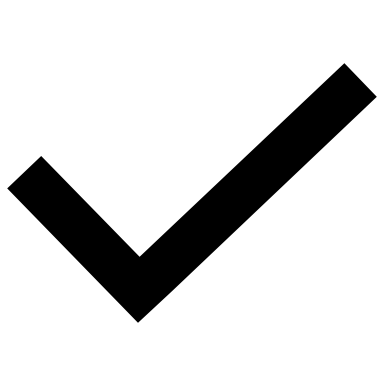 |  | 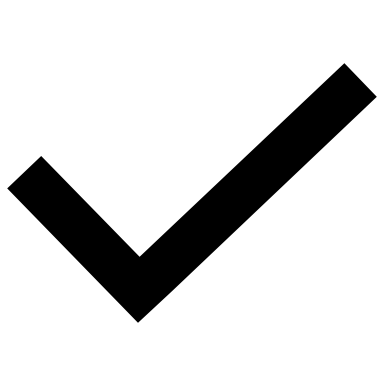 |  |  |  |
| MGI | 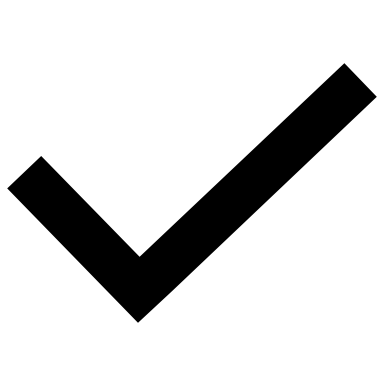 |  | 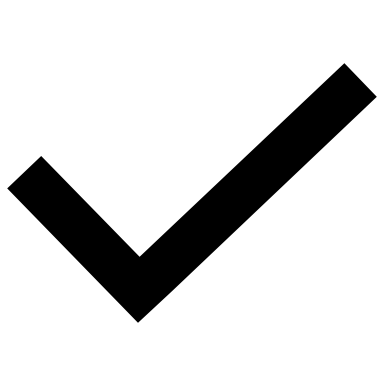 |  |  |  |
| Mouse essential genes | 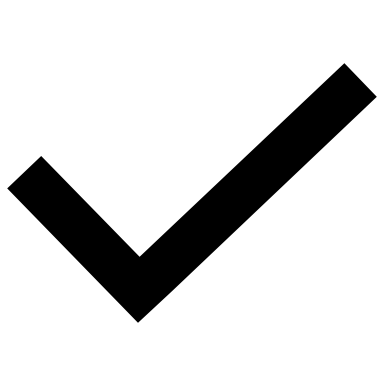 |  | 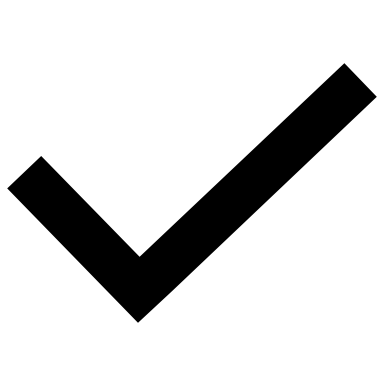 |  |  |  |
| GTEx | 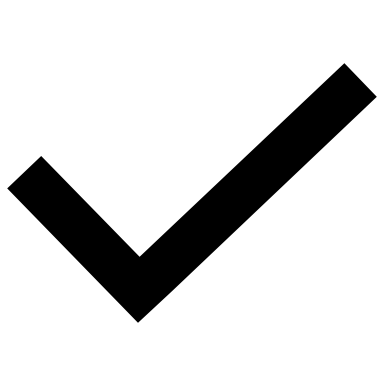 |  | 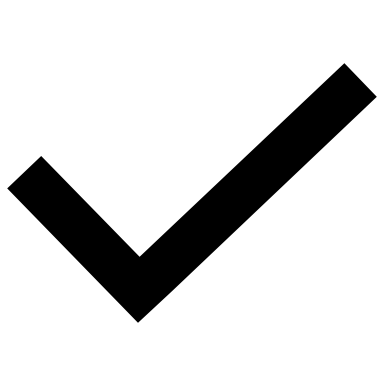 |  |  |  |
| The Human Protein Atlas | 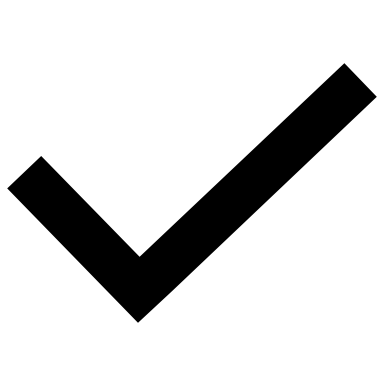 |  | 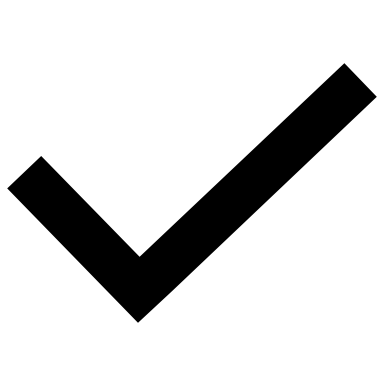 |  |  |  |
| MSigDB | 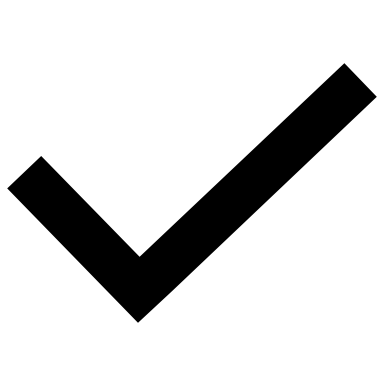 |  | 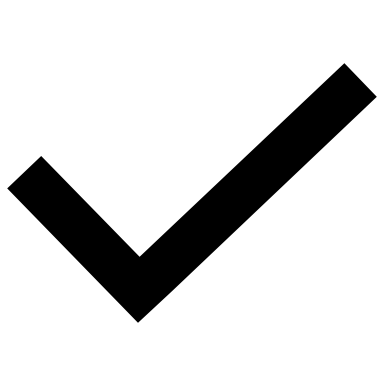 |  |  |  |
| Gene expression (AHBD) |  |  |  | 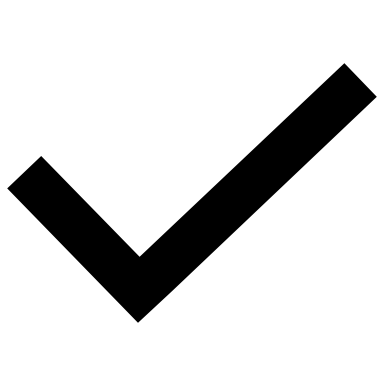 |  | 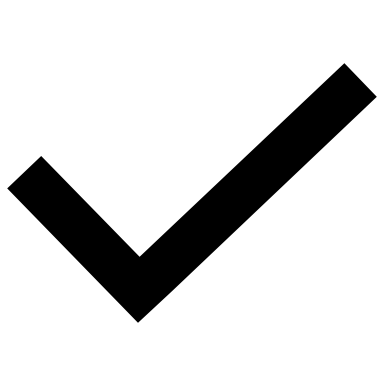 |
| PPI |  |  |  | 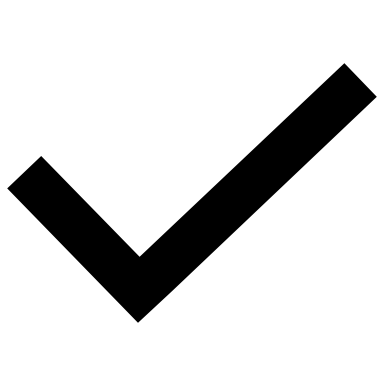 |  | 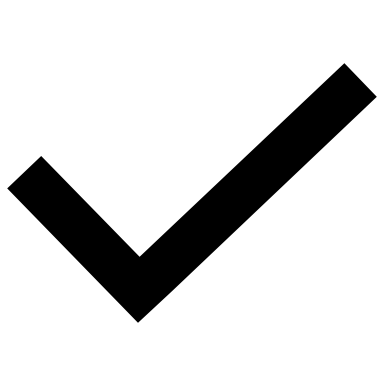 |
| PubMed Literature |  | 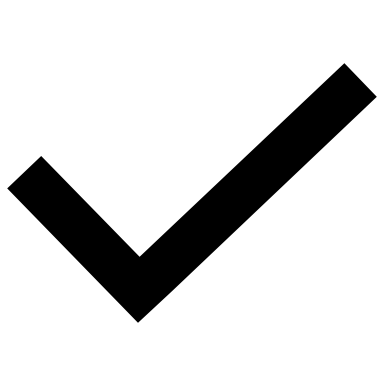 | 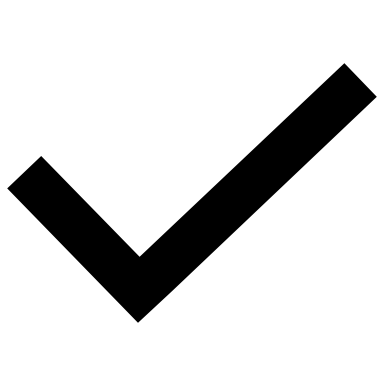 |  | 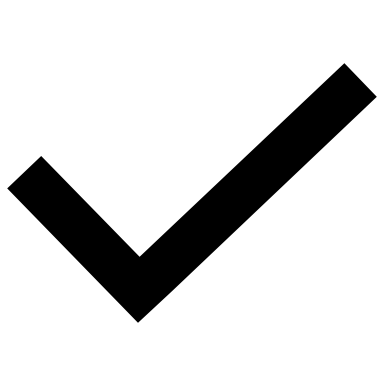 | 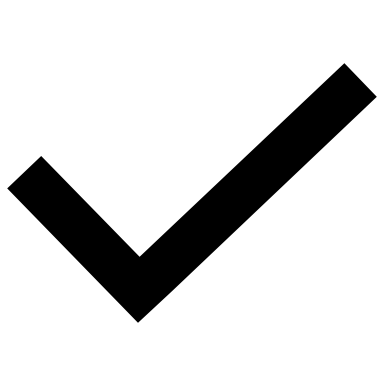 |
| Knowledge Graph (HetioNet) |  |  |  |  | 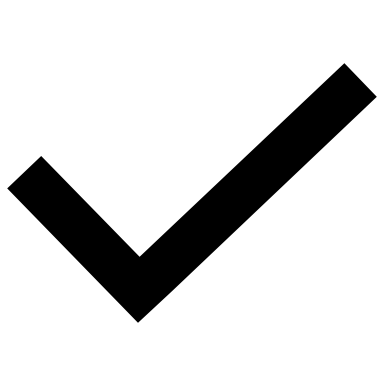 | 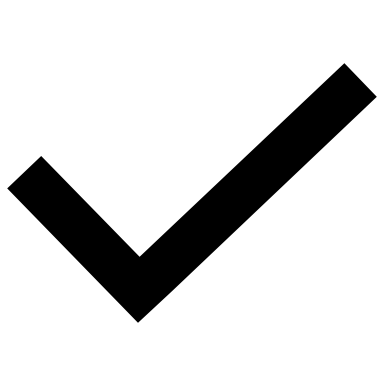 |

Supplementary Table 3. Evaluation results for each method on alternative experimental datasets

| Dataset combination | Median rank obtained through RF (Mantis) framework | Median rank obtained through GNN (SPEOS) framework |
| --- | --- | --- |
| Mantis* | 2281 | 2758 |
| Mantis* + PubMed | 1230 | 1575 |
| Mantis* + PubMed + AHBD gene expression | 1203 | 1432 |
| Mantis* + PubMed + AHBD gene expression + HetioNet KG | N/A | 1419 |
| PubMed | 3109 | 2322 |
| PubMed + AHBD gene expression | 2237 | 1592 |
| PubMed + AHBD gene expression + HetioNet KG | N/A | 1379 |

(Mantis* = All the default datasets used in Mantis framework)

Supplementary Table 4. Precision and Recall rate – temporal-split validation

| precision @ top 100 | | | | | | |
| --- | --- | --- | --- | --- | --- | --- |
| Disease | **RF** | | | **GNN** | | |
|  | **RF** | **C-RF** | **KI-RF** | **GNN** | **C-GNN** | **KI-GNN** |
| DEE | 0.21 | 0.06 | 0.13 | 0.1 | 0.1 | 0.09 |
| AD DEE | 0.12 | 0.07 | 0.08 | 0.06 | 0.02 | 0.08 |
| AR DEE | 0 | 0 | 0.01 | 0 | 0.01 | 0.05 |
| Microcephaly | 0.02 | 0.01 | 0.01 | 0.01 | 0 | 0 |
| Intellectual Disability | 0.15 | 0.01 | 0.07 | 0.05 | 0 | 0.02 |
| Motor Neuron Disease | 0 | 0.01 | 0.02 | 0 | 0.02 | 0.02 |
| Cerebral Palsy | 0.17 | 0.14 | 0.21 | 0.05 | 0.18 | 0.23 |
| Bleeding and Platelet Disorder | 0.07 | 0.06 | 0.07 | 0.01 | 0.04 | 0.07 |
| Arthrogryposis | 0 | 0.05 | 0.03 | 0.00 | 0.03 | 0.04 |

| recall @ 100 | | | | | | |
| --- | --- | --- | --- | --- | --- | --- |
| Disease | **RF** | | | **GNN** | | |
|  | **RF** | **C-RF** | **KI-RF** | **GNN** | **C-GNN** | **KI-GNN** |
| DEE | 0.0805 | 0.023 | 0.0677 | 0.0521 | 0.0521 | 0.0469 |
| AD DEE | 0.1644 | 0.0959 | 0.1096 | 0.0938 | 0.0312 | 0.125 |
| AR DEE | 0 | 0 | 0.0208 | 0 | 0.0217 | 0.1087 |
| Microcephaly | 0.0417 | 0.0208 | 0.0208 | 0.0217 | 0 | 0 |
| Intellectual Disability | 0.0617 | 0.0041 | 0.0288 | 0.0226 | 0 | 0.009 |
| Motor Neuron Disease | 0 | 0.1429 | 0.2857 | 0 | 0.2857 | 0.2857 |
| Cerebral Palsy | 0.0726 | 0.0598 | 0.0897 | 0.0215 | 0.0773 | 0.0987 |
| Bleeding and Platelet Disorder | 0.4375 | 0.375 | 0.4375 | 0.0667 | 0.2667 | 0.4667 |
| Arthrogryposis | 0 | 0.2083 | 0.125 | 0.0 | 0.1364 | 0.1818 |

Supplementary Table 5. Temporal-split validation results on different text datasets and different embedding models using the GNN-based model.

| Text dataset | Embedding model | Other node features | Edges | Median Rank | Mean Rank |
| --- | --- | --- | --- | --- | --- |
| NCBI gene Summary | chatgpt text embedding ada 002 | - | - | 3082 | 4770 |
| NCBI gene Summary | chatgpt text embedding ada 002 | - | KG pathways | 3523 | 4759 |
| NCBI gene Summary | minilm-l6-v2 | - | - | 4361 | 5112 |
| NCBI gene Summary | minilm-l6-v2 | - | KG pathways | 3262 | 4441 |
| PubMed Titles | minilm-l6-v2 | - | - | 1503 | 3280 |
| PubMed Titles | minilm-l6-v2 | - | KG pathways | 1649 | 3429 |
| PubMed Titles | chatgpt text embedding ada 002 | - | - | 2108 | 4190 |
| PubMed Titles | chatgpt text embedding ada 002 | - | KG pathways | 2016 | 3963 |
| PubMed Titles and Abstracts | chatgpt text embedding ada 002 | - | - | 2108 | 4190 |
| PubMed Titles and Abstracts | chatgpt text embedding ada 002 | - | KG pathways | 2016 | 3963 |
| PubMed titles | chatgpt text embedding ada 002 | AHBD expression | - | 1275 | 2566 |
| PubMed titles | chatgpt text embedding ada 002 | AHBD expression | KG pathways | 1426 | 2727 |
| PubMed titles | minilm-l6-v2 | AHBD expression | - | 1433 | 2748 |
| PubMed titles | minilm-l6-v2 | AHBD expression | KG pathways | 1348 | 2615 |

Supplementary Table 6. Comparison of different knowledge graph edge types under temporal-split evaluation (Lower is better.). Node features: gene expression + PubMed embeddings.

| Edge type (knowledge graph connection) | Median | Mean |
| --- | --- | --- |
| Pathway | **1379** | **2616** |
| Co-variation | 1833 | 3380 |
| Interaction | 1744 | 2943 |
| Regulation | 1496 | 2946 |
| Biological Process (1/3 subsample) | 2134 | 3239 |
| Neural Biological Process | 1544 | 2929 |
| Disease | 1426 | 3112 |
| Cellular Component | 1549 | 2972 |
| Molecular Function | 1800 | 2999 |
| Compound | 1784 | 3367 |
| Pathway + Neural Biological Proces | 1418 | 2934 |
| Pathway + Disease | 1426 | 2891 |
| Pathway + Regulation | 1401 | 2747 |
| Pathway + Disease + Regulation | 1403 | 2899 |

**Supplementary Figures**

Biological Evaluation Results for AD DEE genes

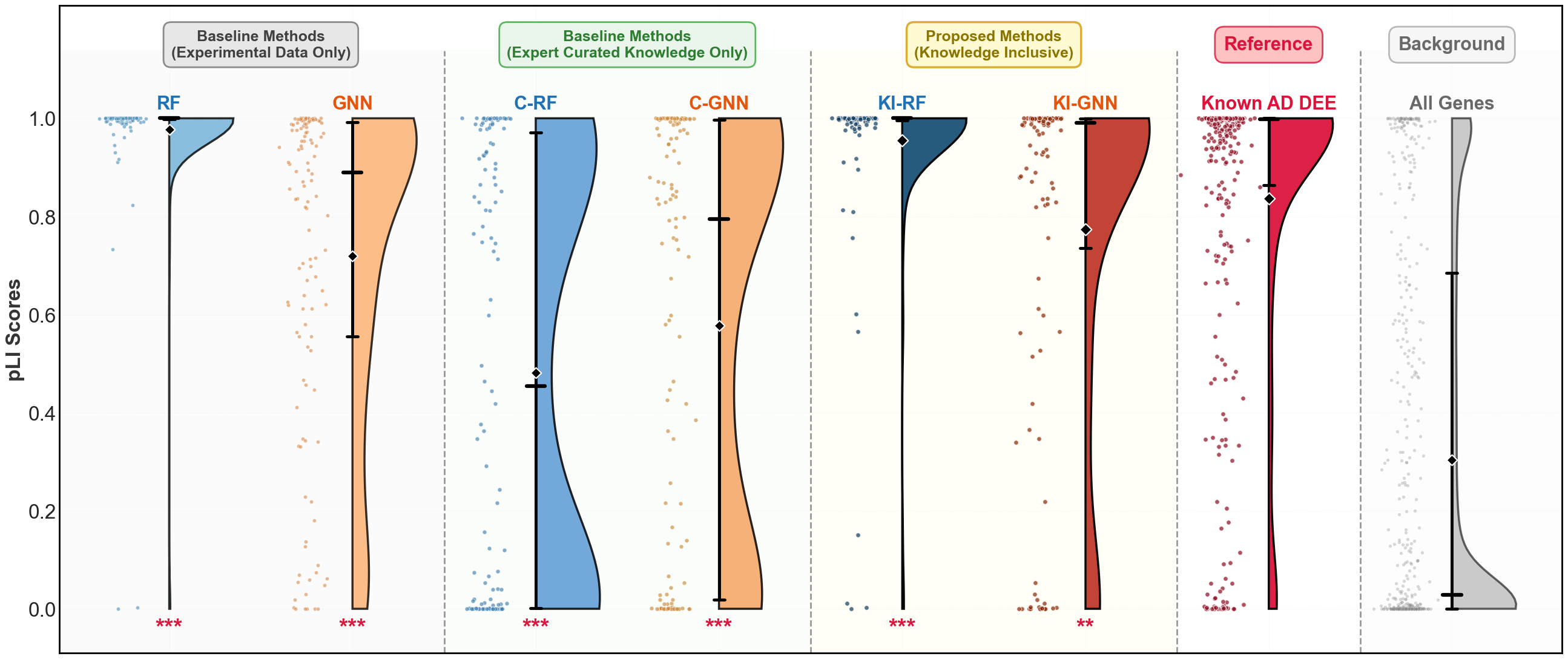

Supplementary Fig. 1: Distribution of probabilities of LoF intolerance (pLI) scores for AD DEE candidate genes predicted by different methods, along with the distribution of known AD DEE genes (reference, in red) and all human genes (background, in grey). Existing methods (RF, GNN) are shown on the left, expert-curated knowledge only methods (C-RF, C-GNN) in the middle, and proposed Knowledge Inclusive methods (KI-RF, KI-GNN) on the right. The distributions were compared against known DEE genes using the Mann-Whitney U test, with significance levels indicated below each method (significance levels: *** p<0.001, ** p<0.01, * p<0.05, ns = not significant).

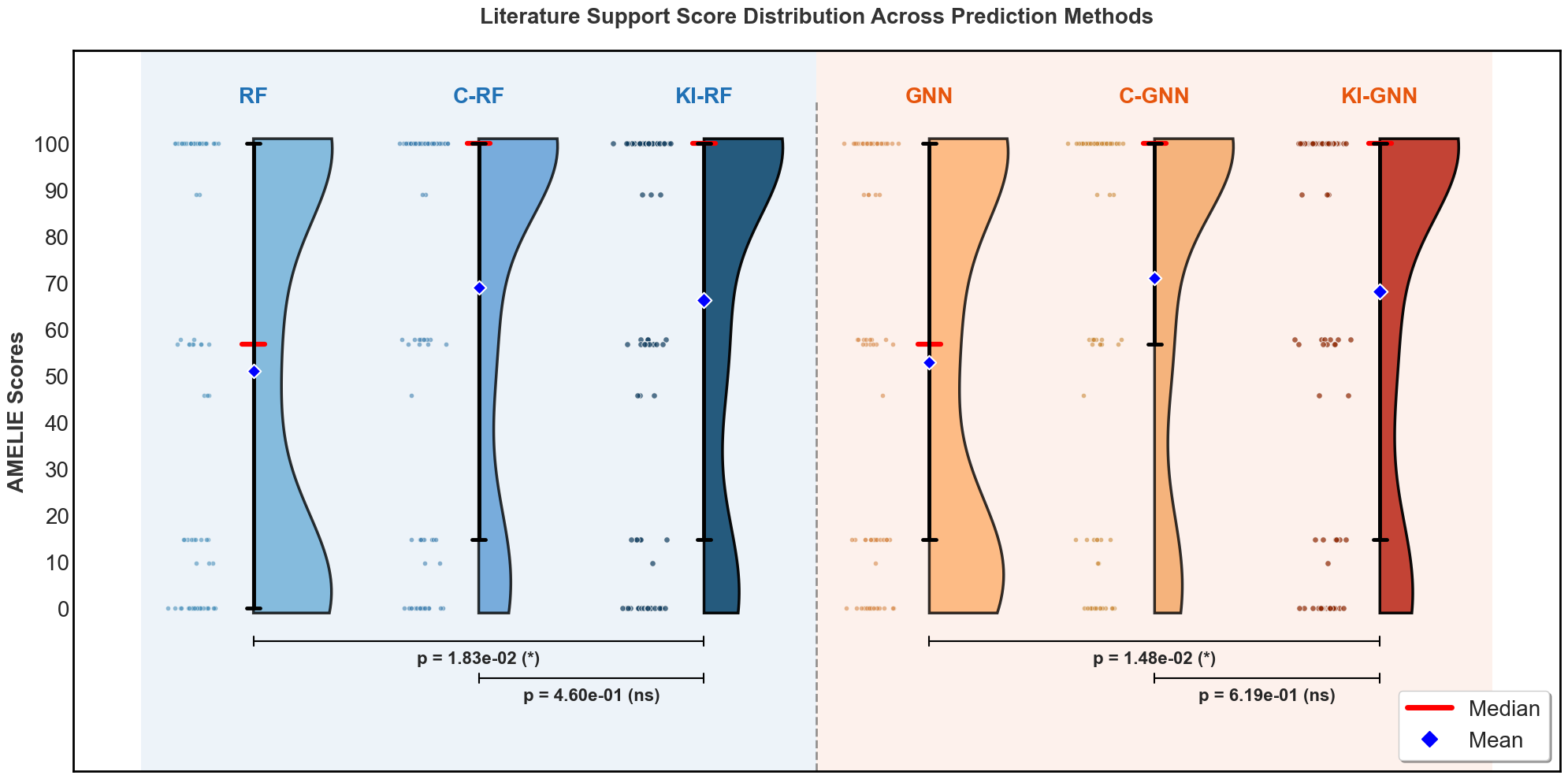

Supplementary Fig. 2: AMELIE scores for the top 100-ranked AD DEE candidate genes across different methods. The red line indicates the median score, and the blue diamond marker indicates the mean score. Pairwise comparisons were performed using the Mann–Whitney U test, with corresponding p-values shown above the plot (significance levels: *** p*<*0.001, ** p*<*0.01, * p*<*0.05, ns = not significant).

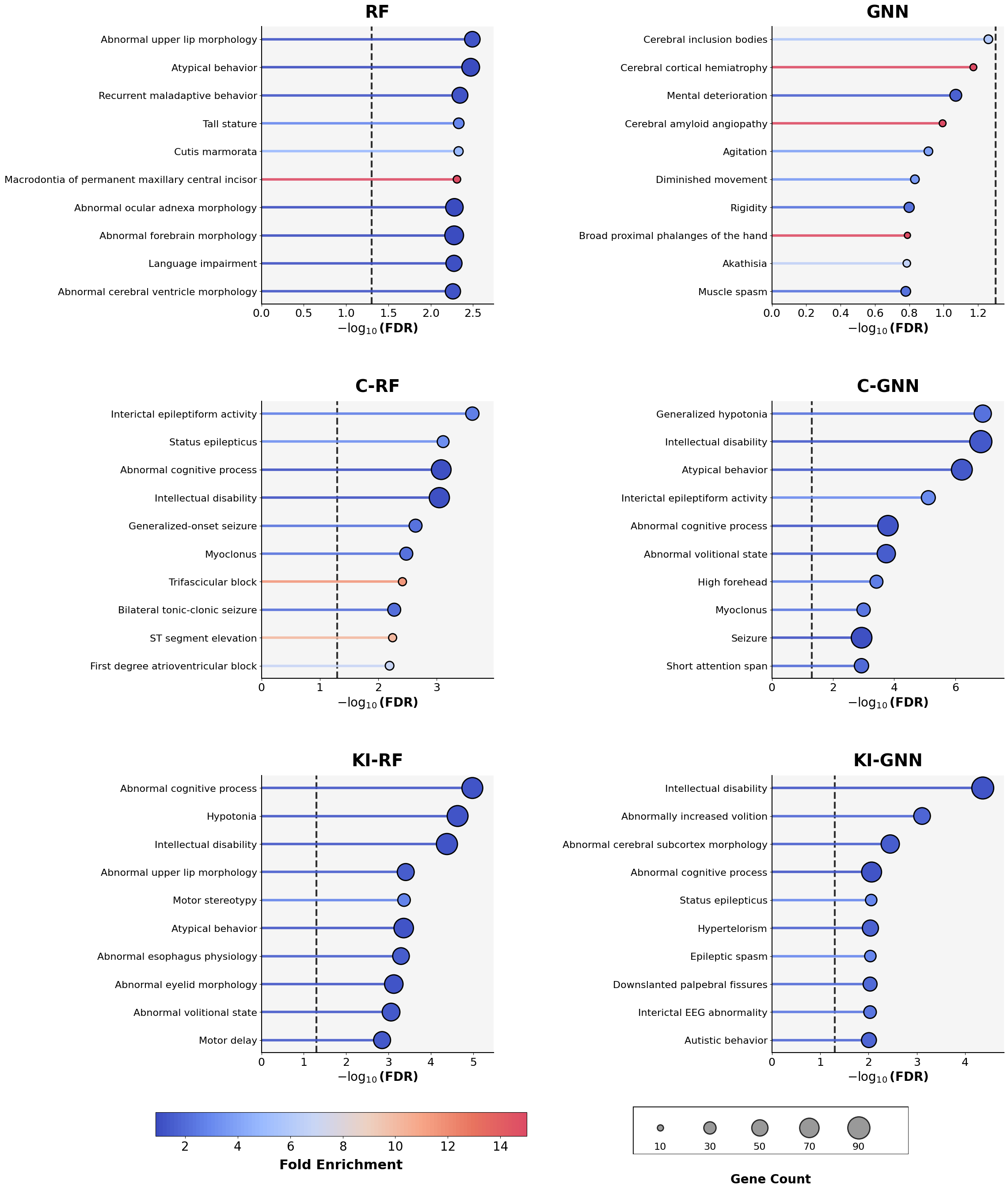

Supplementary Fig. 3: Phenotype enrichment evaluation results for the top 100 AD DEE candidate genes. The top 10 enriched phenotype terms from each method are shown as horizontal bar plots. Bar lengths represent the statistical significance of enrichment (-log (FDR)), with the vertical dashed line indicating the FDR cutoff (FDR = 0.05; -log (FDR) = 1.3). Circle sizes at the bar ends denote the number of candidate genes (out of 100) associated with each term, while colours indicate the fold enrichment values.

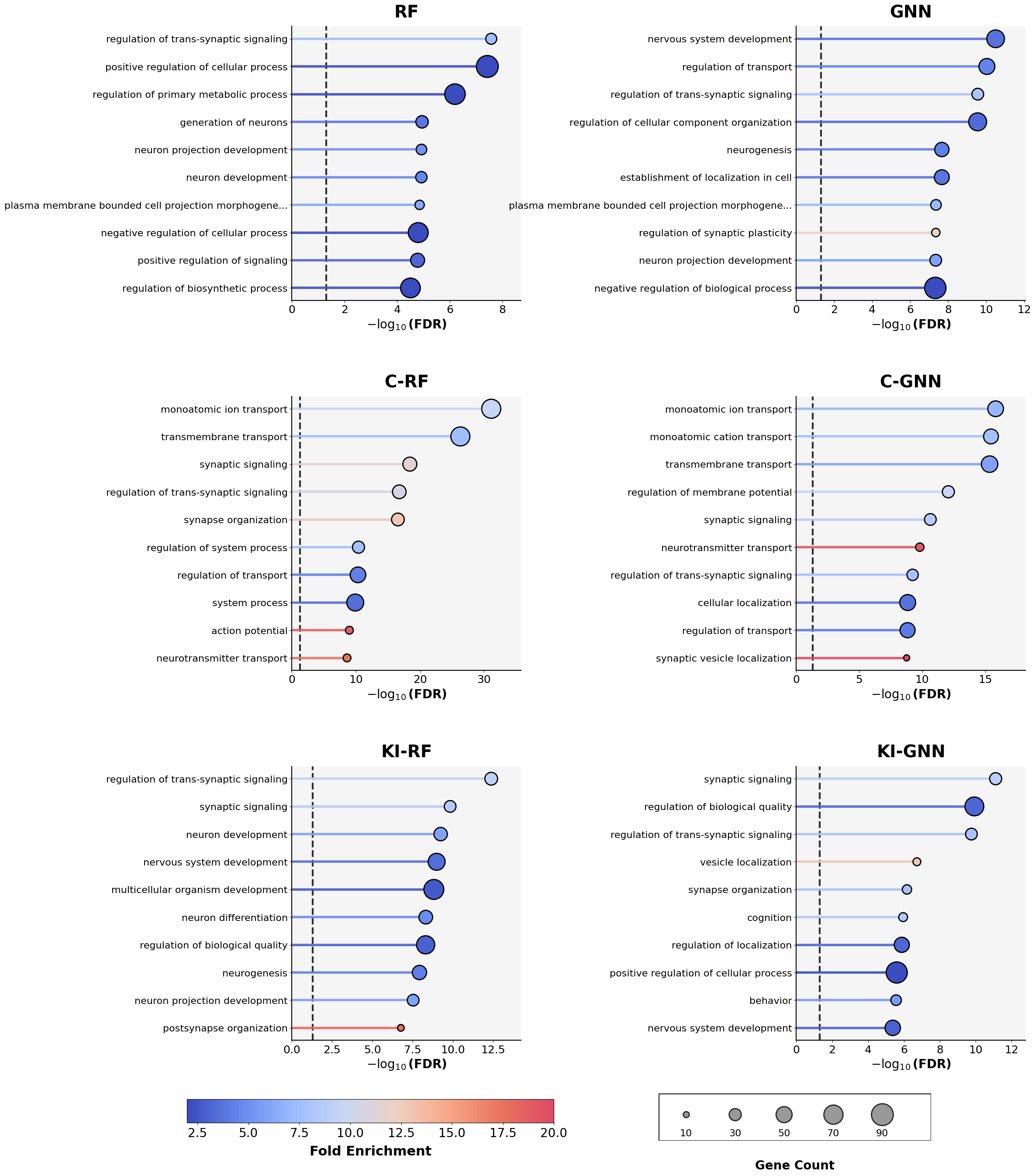

Supplementary Fig. 4: Gene Ontology enrichment evaluation results for the top 100 AD DEE candidate genes. The top 10 enriched Gene Ontology (GO) terms from each method are shown as horizontal bar plots. Bar lengths represent the statistical significance of enrichment (-log (FDR)), with the vertical dashed line indicating the FDR cutoff (FDR = 0.05; -log (FDR) = 1.3). Circle sizes at the bar ends denote the number of candidate genes (out of 100) associated with each GO term, while colours indicate the fold enrichment values.

Biological Evaluation Results for AR DEE genes

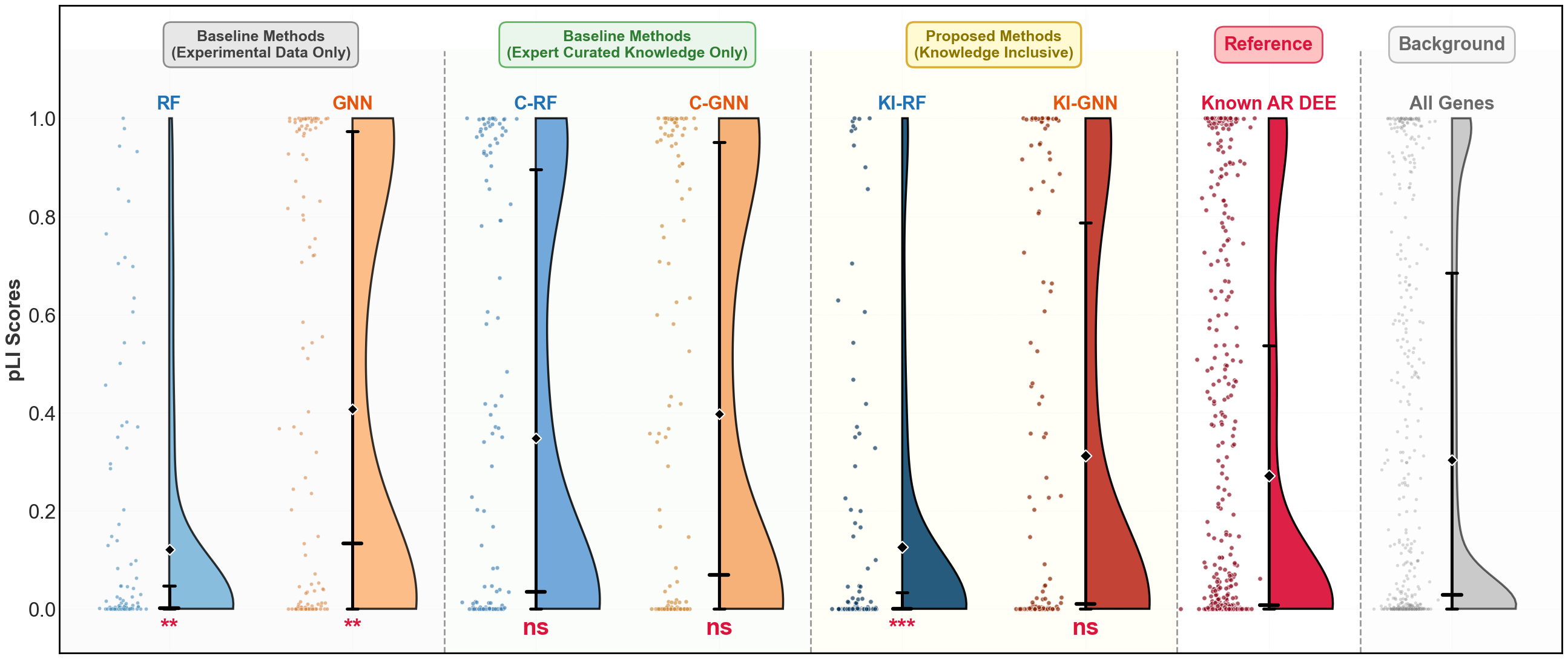

Supplementary Fig. 1: Distribution of probabilities of LoF intolerance (pLI) scores for AR DEE candidate genes predicted by different methods, along with the distribution of known AR DEE genes (reference, in red) and all human genes (background, in grey). Existing methods (RF, GNN) are shown on the left, expert-curated knowledge only methods (C-RF, C-GNN) in the middle, and proposed Knowledge Inclusive methods (KI-RF, KI-GNN) on the right. The distributions were compared against known DEE genes using the Mann-Whitney U test, with significance levels indicated below each method (significance levels: *** p<0.001, ** p<0.01, * p<0.05, ns = not significant).

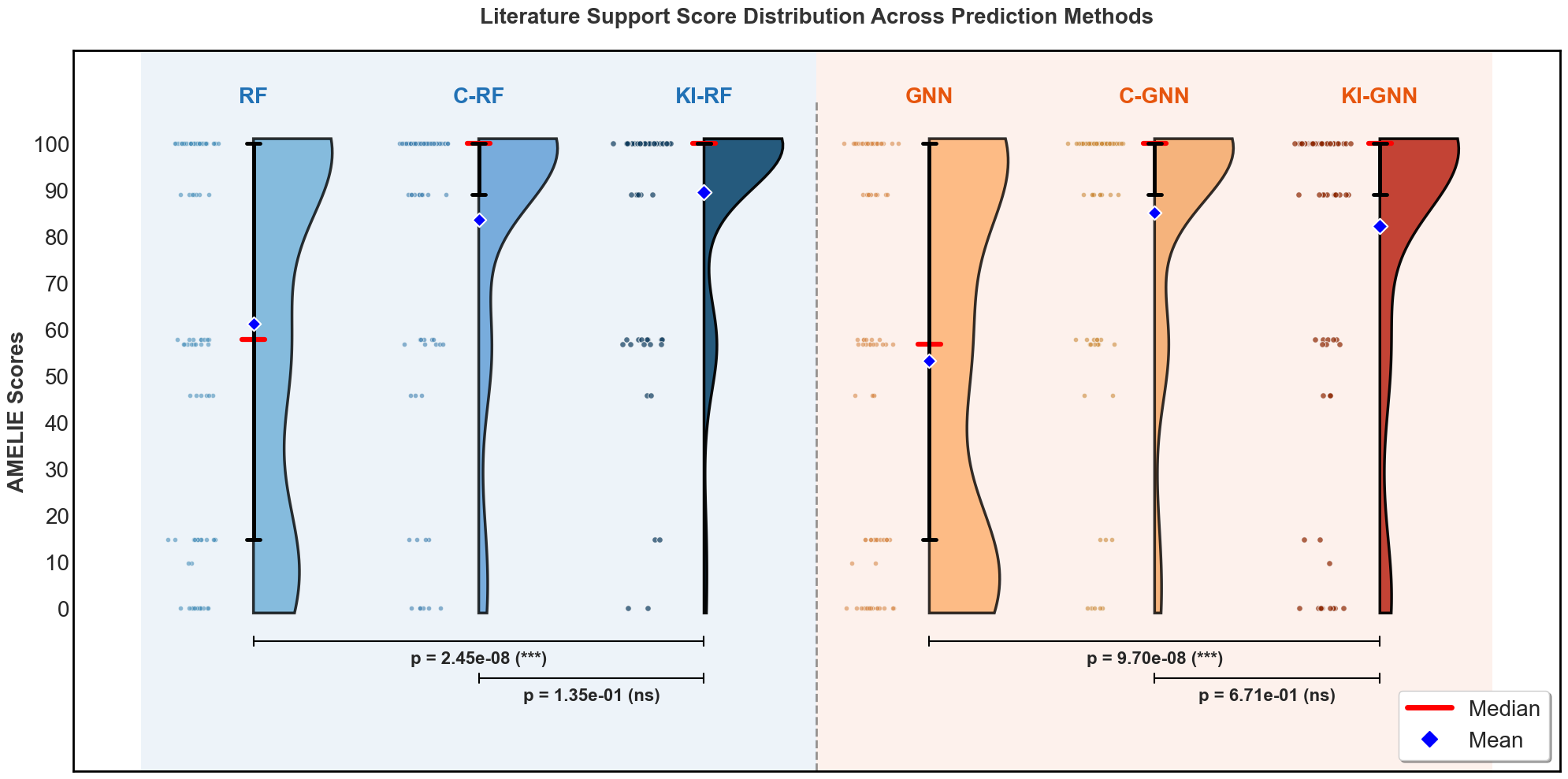

Supplementary Fig. 6: AMELIE scores for the top 100-ranked AR DEE candidate genes across different methods. The red line indicates the median score, and the blue diamond marker indicates the mean score. Pairwise comparisons were performed using the Mann–Whitney U test, with corresponding p-values shown above the plot (significance levels: *** p*<*0.001, ** p*<*0.01, * p*<*0.05, ns = not significant).

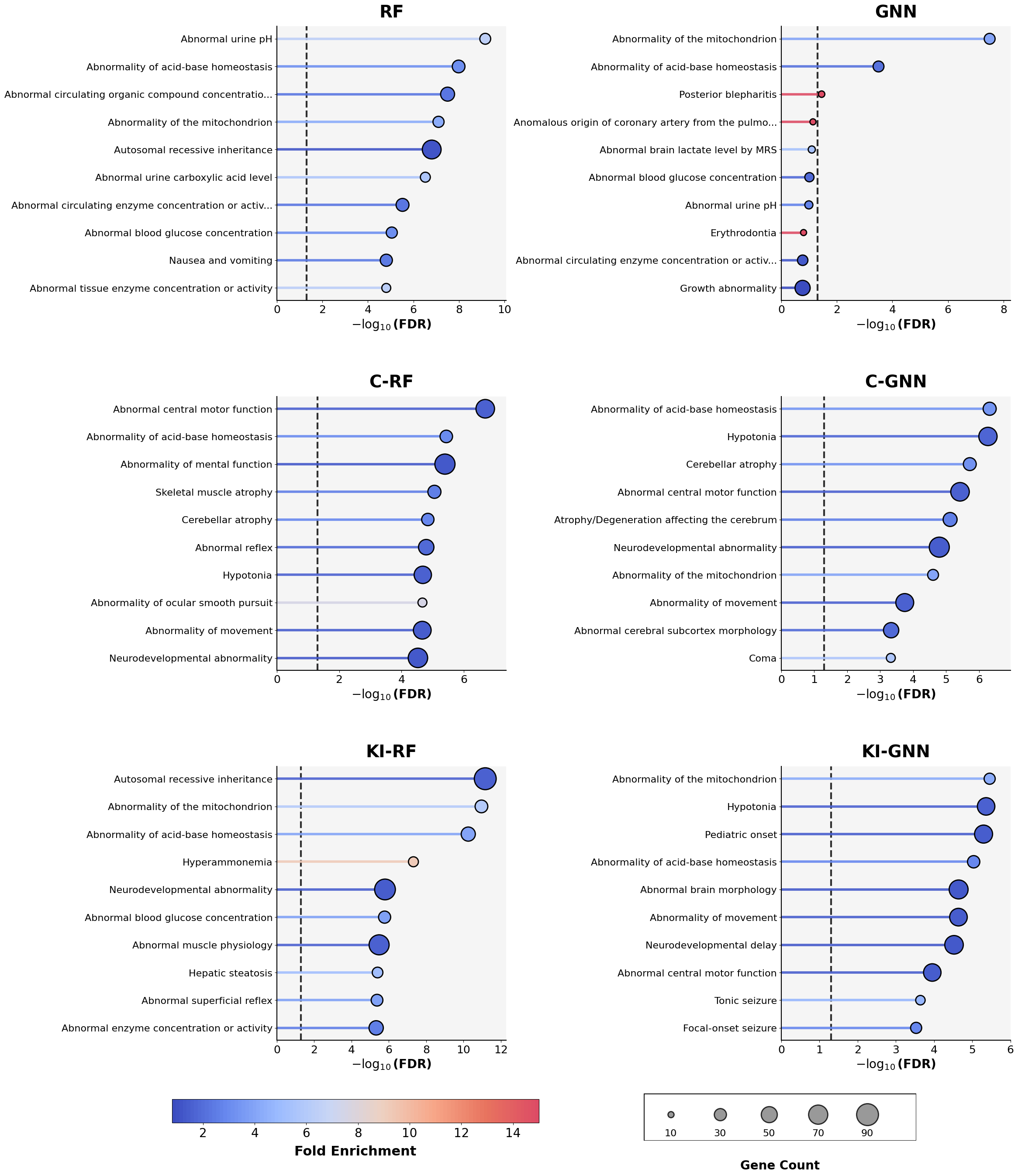

Supplementary Fig. 7: Phenotype enrichment evaluation results for the top 100 AR DEE candidate genes. The top 10 enriched phenotype terms from each method are shown as horizontal bar plots. Bar lengths represent the statistical significance of enrichment (-log (FDR)), with the vertical dashed line indicating the FDR cutoff (FDR = 0.05; -log (FDR) = 1.3). Circle sizes at the bar ends denote the number of candidate genes (out of 100) associated with each term, while colours indicate the fold enrichment values.

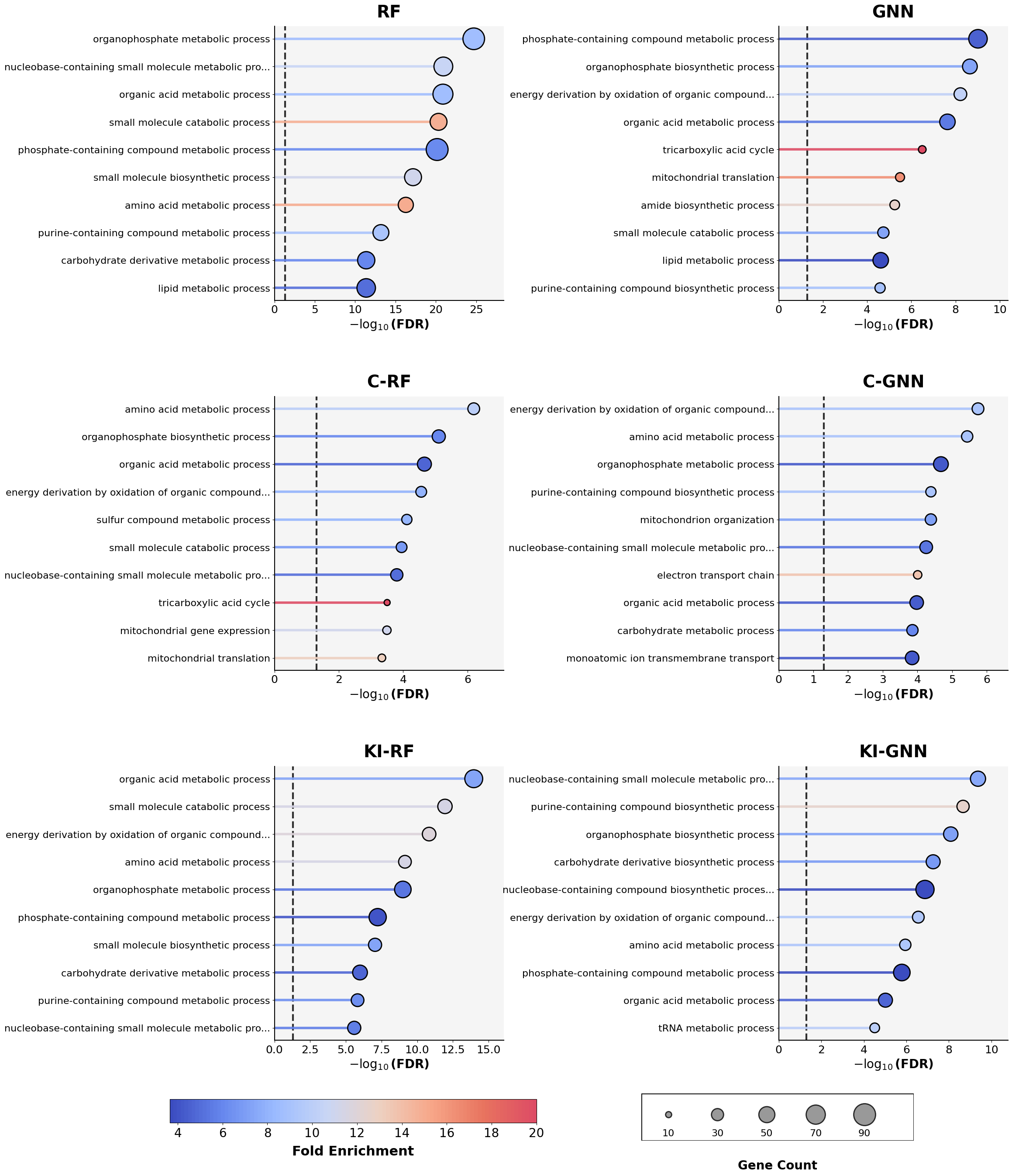

Supplementary Fig. 8: Gene Ontology enrichment evaluation results for the top 100 AR DEE candidate genes. The top 10 enriched Gene Ontology (GO) terms from each method are shown as horizontal bar plots. Bar lengths represent the statistical significance of enrichment (-log (FDR)), with the vertical dashed line indicating the FDR cutoff (FDR = 0.05; -log (FDR) = 1.3). Circle sizes at the bar ends denote the number of candidate genes (out of 100) associated with each GO term, while colours indicate the fold enrichment values.

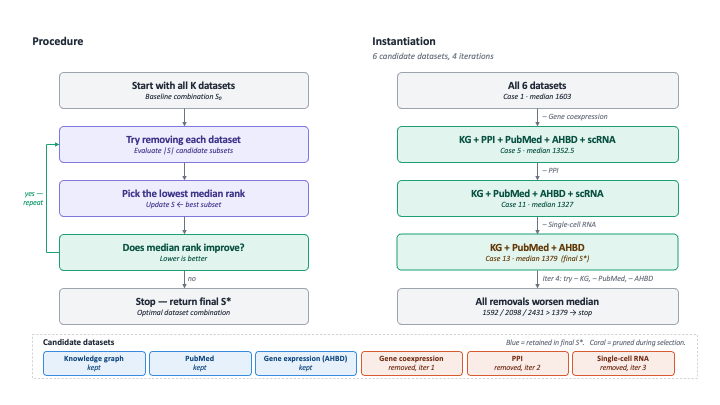
 Supplementary Fig 9: Ablation study identifying the optimal dataset combination. At each iteration, one dataset is removed from the current best subset; the removal producing the lowest median rank is retained. Detailed results for all cases are in Supplementary Data 1.
